# Nanodisc, amphipol or detergent belts in cryoEM reconstructions of membrane proteins are similar and correspond to a common ordered solvent layer

**DOI:** 10.1101/2020.12.10.418871

**Authors:** Veronica Zampieri, Alexia Gobet, Xavier Robert, Pierre Falson, Vincent Chaptal

## Abstract

To maintain membrane proteins soluble in aqueous solution, amphipathic compounds are used to shield the hydrophobic patch of their membrane insertion, which forms a belt around the protein. This hydrophobic belt is seldom looked at due to the difficulty to visualize it. Cryo-EM is now offering this possibility, where belts are visible in 3D reconstructions. We investigated membrane proteins solved in nanodiscs, amphipols or detergents to analyze whether the nature of the amphipathic compound influences the belt size in 3D reconstructions. We identified belt boundaries in map-density distributions and measured distances for every reconstruction. We showed that all the belts create on average similar reconstructions, whether they originate from the same protein, or from protein from different shapes and structures. There is no difference among detergents or types of nanodisc used. These observations illustrate that the belt observed in 3D reconstructions corresponds to the minimum ordered layer around membrane proteins.

## 1. Introduction

Membrane protein structure determination has become almost a routine job with the recent development of single particle electron microscopy in cryogenic conditions (Cryo-EM). A skyrocketing amount of membrane protein structures becomes available improving our knowledge of many biological processes. In order to achieve this grail of a nice quality structure, it is necessary to extract the protein from the native membrane, and purify it to homogeneity so it can be applied on a grid and imaged on a microscope. And there lies the specificity of membrane proteins: they display a part of their structure that spans the membrane, abundant in hydrophobic residues, rendering them insoluble in water. There is thus a need for some amphipathic compound to shield this trans-membrane region from water and from other hydrophobic molecule or even other proteins around, else the result will be aggregation and loss of the precious gem.

Many recipes are available today to maintain membrane proteins in solution. The historical way, still very much used today, is to use detergents to extract membrane proteins from the membrane and then purify them in detergent solutions. Detergents are small molecules that display a hydrophilic head and a hydrophobic tail. Both moieties vary in nature, length and size allowing a large panel of possible screening for good conditions, and they are also sometimes used in mixtures[1]. By nature, detergents are very mobile and form a dynamic belt wrapping around the trans-membrane part of the protein[2]. Due to this dynamic property, detergents can have sometimes negative impacts on membrane proteins structure and function. Therefore, detergents with increased stabilizing properties have been more recently conceived for limiting such mobility either by having a design close to lipids (LMNG)[3] or by generating specific interactions[4]. Also, their amphipathic nature is unique to stabilize given conformations. Other tools have been developed to forgo the need for detergents. Among them the derivation of the lipid A apolipoprotein engineered as a series of Membrane Scaffold Proteins (MSP), that together with lipids and the membrane protein will form a lipidic nanodisc is a real success, allowing to reconstitute a more native environment and/or to vary the type of lipids around the membrane protein[5]. In the same vein, amphipols are polymers that wrap around purified membrane proteins and stabilize them without the need for detergents and lipids[6]. All these tools have been used for membrane protein structure determination by Cryo-EM. More recently, new polymers have been designed to directly extract membrane proteins from native membranes, allowing their purification without detergents[7, 8].

All these compounds generate a local amphipathic environment around the membrane region of membrane proteins that maintains them in aqueous solutions. This layer is a belt from which membrane proteins are indissociable. This solvent is however seldom looked at despite its huge influence on the protein function, due to the difficulty to visualize it. Cryo-EM now allows for the visualization of a layer wrapping around the transmembrane region of membrane proteins. We have taken this opportunity to investigate if there is an influence of these different amphipathic belts on their visualization after 3D reconstructions. We identified the position of the hydrophobic solvent belt in map-density distributions and measured the belts for many different proteins solved in nanodiscs, amphipols and detergents. We showed that all the belts create on average similar reconstructions, whether they originate from the same protein, or from protein from different shapes and structures. There is no difference amongst detergents or type of nanodisc used. These observations illustrate that the averaging procedure of CryoEM reconstructions returns a belt corresponding to a common minimum ordered solvent layer amongst all the membrane proteins particles, selected for this reconstruction. This solvation layer is however much smaller than the total amount of amphipathic compound embarked around membrane proteins.

## 2. Results

### 2.1. 3D reconstructions of membrane proteins structures solved by cryo-EM in several amphipathic belts

In order to discriminate whether there is an influence of the type of amphipathic belt on the 3D reconstructions used to image membrane proteins by cryo-EM, we have screened the whole Protein Data Bank to select for comparison the membrane proteins that have been solved only in multiple hydrophobic environments: nanodiscs, amphipols or detergents (Fig. 1A. and STable 1.). The idea behind this selection is to keep the protein fold constant in order to normalize its influence on the reconstruction, and to be able to focus on the solvent belts alone. The identification of the belt is obvious to a trained eye, capable of detecting the trans-membrane parts of a protein in a structure. The hydrophobic belt is characterized by an expansion of lower-level density in the vicinity of the membrane region as a decrease in the map density. After the observation of map-density distributions of each structure of this dataset, an apparent feature was observable to identify the belt and is exemplified in Figure 1B. At high density levels, the very ordered parts of the structure are visible, on which reconstruction was anchored. Typically, trans-membrane helices are key features used in 3D reconstructions of membrane proteins and are visible at this level. With decreasing density levels, the number of voxels increases in a concave shape (level 0). The higher ordered layers of the belt start to appear when the curve becomes convex (level 1, dotted arrow). The belt becomes more and more apparent over the course of about one log when the curve inflexes concavely (level 2) before a sharp increase in number of voxels leading to appearance of low-level noise throughout the box (level 3). Across all reconstructions, it is apparent that the more visible the hydrophobic belt, the clearer and sharper the transition is between levels 1 and 2.

**Fig. 1.**
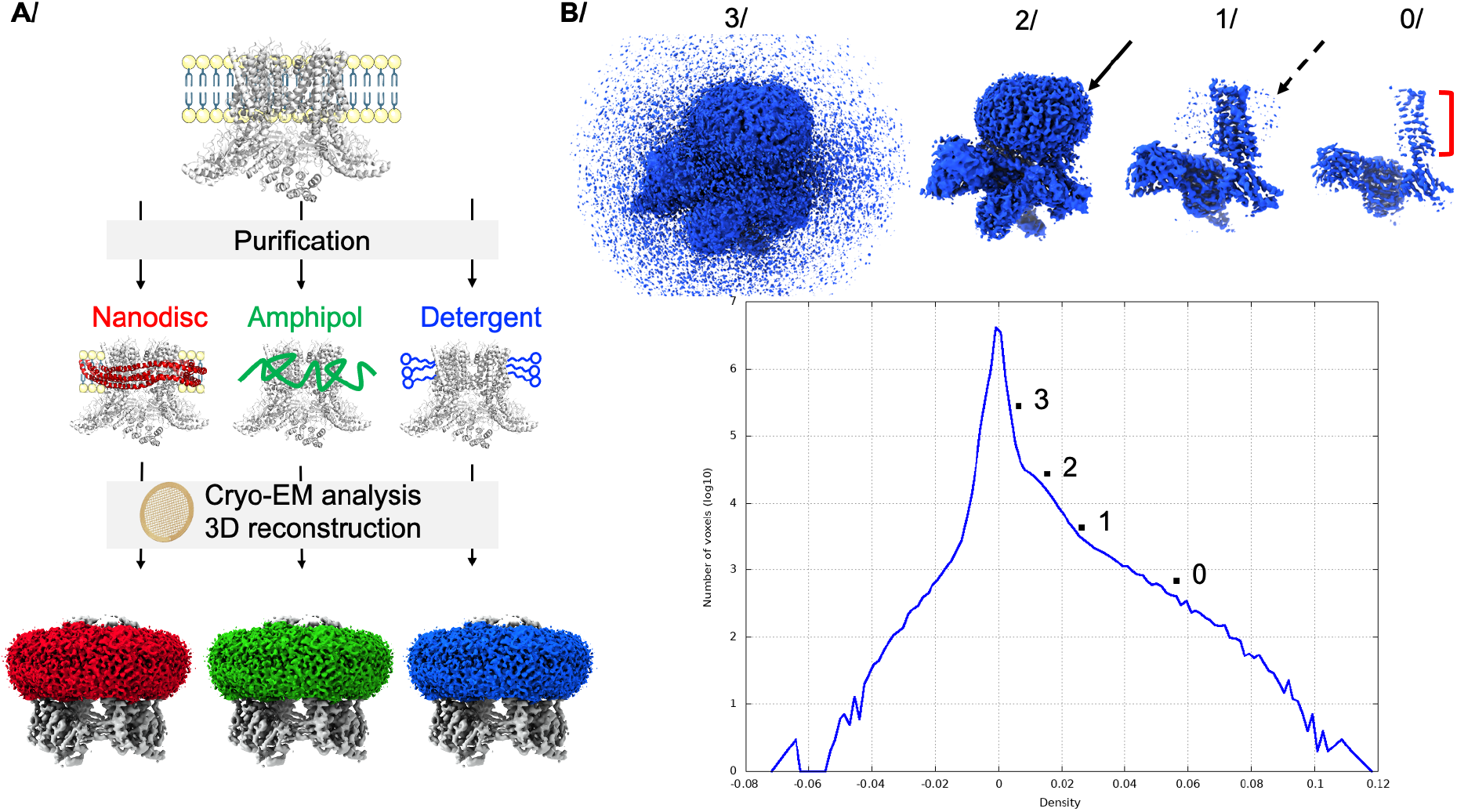
Visualization of the different types of hydrophobic belts surrounding membrane proteins. **A/** General scheme of membrane protein purification from the membrane, kept in detergent (blue) or reconstituted in nanodisc (red) or amphipol (green), and imaged by cryo-EM to obtain a 3D reconstruction. The channel TRPV1 is used as an example of reconstruction (EMD-8117), with belts colored accordingly. **B/** Typical map-density distribution and representative density levels of reconstructions (EMD-20079). Level 0 corresponds to parts of the structure with strong density; the red bar shows the trans-membrane domain. Level 1 corresponds to the appearance of high density for the belt, depicted by the dotted arrow. Level 2 represents the maximum density observed at low-level density, depicted by the solid arrow. Level 3 corresponds to low density noise.

### 2.2. 3D reconstructions of membrane proteins in nanodiscs or detergents yield similar average belt sizes

Using the map-density histogram, we measured for each entry the belt size by determining the distance between the protein edge and the solvent boundaries at level 2 (SFig. 2 to 16., STable 2.). Belt reconstructions are not spherical but rather follow the protein shape. We could thus identify large and small distances of the hydrophobic solvent belt, which we separated in two categories for further processing. Figure 2 displays the distance distribution plot of all proteins separated by hydrophobic environments: nanodiscs, amphipols and detergents. For each type of belt, there is an apparent spread of distances, with 95% of total distances comprised between 14 and 36 Å, and no distances bellow 10 Å around the protein. Statistical analysis of long distances observed in detergents and nanodiscs show that they follow the same distribution, as well as small distances for these two categories. On average, the solvent belt is visible around the protein for 21 to 27 Å. The smaller amount of structures solved using amphipols precludes the statistical analysis on means using parametric statistics. Non-parametric statistics on ranks reveals first an ambiguity about long amphipol distances where the current data set cannot distinguish whether the distances are different or similar (more measurements on more structures are needed to solve the debate), and second unambiguously state that short distances measured in amphipols follow the same distribution as the nanodiscs or detergents ones. Put together, these results point to a common average distance distribution of solvent belts surrounding membrane protein observed after 3D reconstruction of cryo-EM data.

**Fig. 2.**
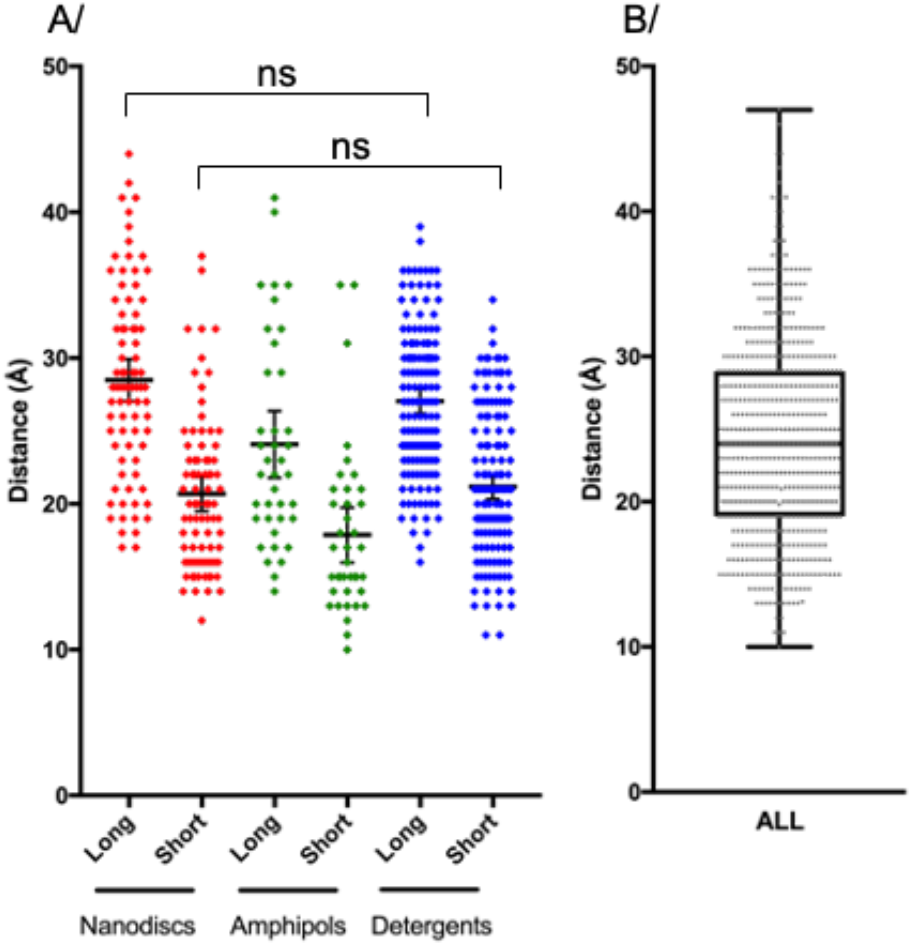
Distance distributions between the protein edge and the belt boundary. **A/** Distances separeted by type of solvent. Nanodiscs (red), amphipols (green) and detergents (blue). Each dot corresponds to a distance, the horizontal bar is the mean with the error bar dispalying the 95%, confidence interval of the mean (reported in, brackets bellow). Long and short distances are, separated for clarity. Nanodiscs, long 28 Å [27–30], and short 21 Å [20–22]. Amphipols, long 24 Å [22–26] and short 18 Å [16–20]. Detergents, long 27 Å [26–28] and short 21 Å [20–22]. **B/** All distances represented by a grey dot. The box corresponds to the 25^th^ (19 Å) and 75^th^ (29 Å) percentile with an equal mean and median of 24 Å. 95% of total, distances are comprised between 14 and 36 Å.

In order to distinguish if there is some inter-family of inter-solvent specificities hidden within the global distribution, we further separated proteins for individual analysis.

### 2.3. The TRPV family

The Transient Receptor Potential Vanilloid (TRPV) family consists of six ion channels, varying in ion-selectivity according to the sub-family. Despite being functionally distinct, they share a highly conserved fold, being active as a tetramer formed around 6 trans-membrane helices per monomer[9]. In the present dataset, we identified structures of TRPV1 in nanodiscs and amphipols, TRPV2 in nanodiscs, amphipols and detergents (LMNG and DMNG), and TRPV5 in nanodiscs and DMNG (STable 1.). The fact that TRPV proteins share a conserved fold gives a unique opportunity to compare varying hydrophobic solvent belts. To these proteins, we also included structures of TRPV3 and 6 that were solved in only one type of amphipathic solvent, benefiting from the fact that they share the same fold.

The signal of the hydrophobic belt varies in intensity among the different 3D reconstructions, for unclear reasons (Fig. 3A., SFig. 2–5.). For example, the nanodisc belt of TRPV1 and 2 appears with a strong signal in these five reconstructions, while its intensity is much milder in the two reconstructions of TRPV5. Similar trends can be seen in amphipols or detergents across the various reconstructions. Nevertheless, belt boundaries are clearly visible and were measured for all thirty proteins (Fig. 3C.) Distance distributions follow a similar trend as the global one (Fig. 2A.), where differences amongst belts are undistinguishable. The same ambiguity remains between long distances of nanodiscs and amphipols, but it is challenged by the lack of difference this time between amphipols and detergents. More measures on more reconstructions would help to differentiate the trend. Nevertheless, the fact that short distances show the same distributions across the three types of solvent and the undistinguishable long distances for nanodiscs and detergents suggest a similar average size for solvent belts around TRPV proteins.

**Fig. 3.**
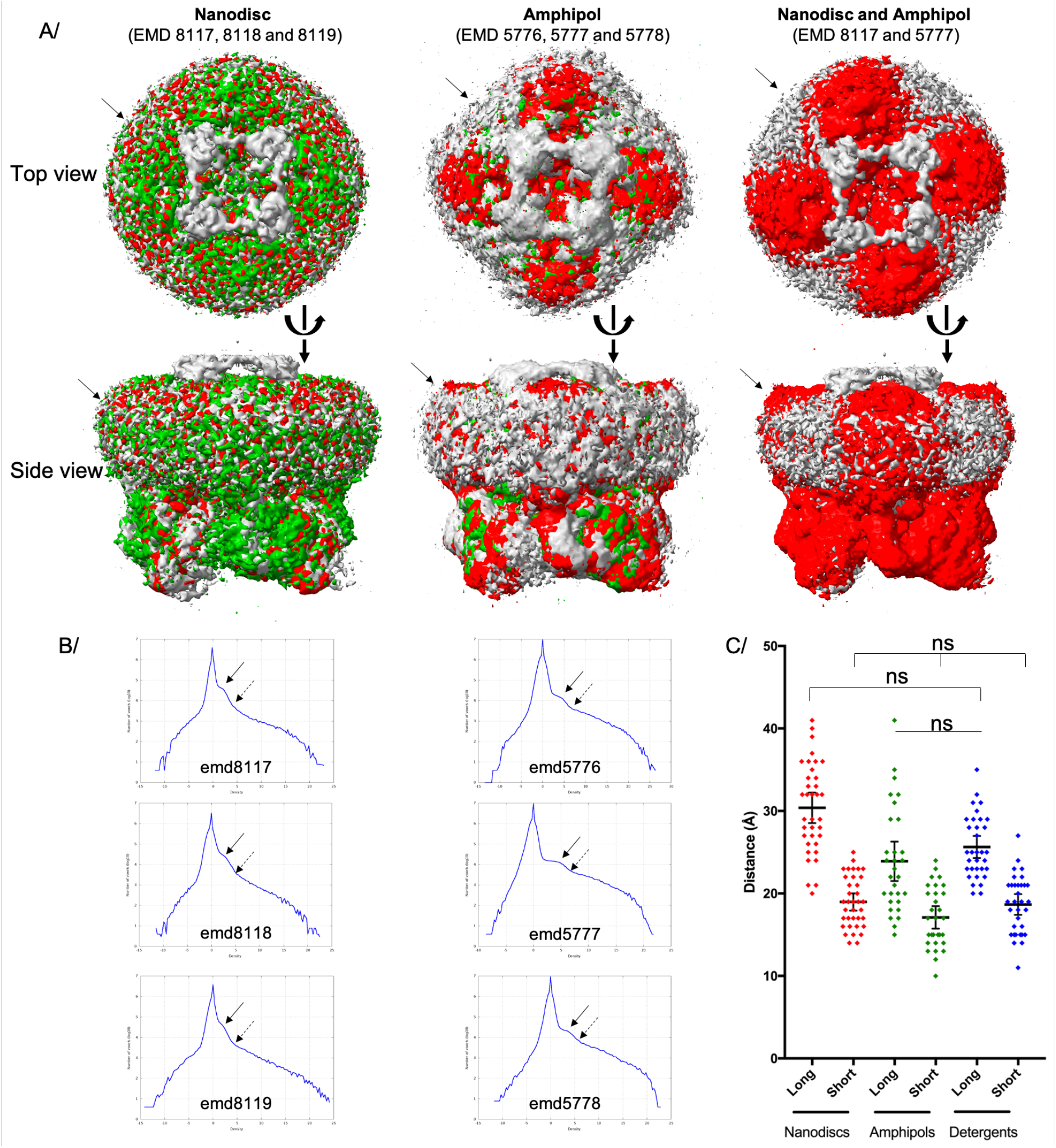
Belts surrounding TRPV proteins. **A/** Example of the TRPV1 proteins solved in nanodisc or amphipols. The EMD accession codes are listed and each structure are colored in grey, red or green, and overalaid. Top and side views are depicted, the solid arrow points to the position of the solvent belt. **B/** Map-density distribution for each entry. On each distribution, the dotted arrow shows the belt appearance at level 1, and the solid arrow at level 2. **C/** Distance distributions for all TRPV proteins, with the same color-coding as Fig.2. Statistical analysis was carried out using ANOVA and Kruskal-Wallis.

### 2.4. Similar average belts length across multiple protein types solved using different amphipathic solvents

We have identified in the dataset multiple protein structures that have been solved only a few times in different hydrophobic belts. The limited amount of structures prevents a statistical analysis on each protein. Instead, these proteins were evaluated in a group, thereby offering the opportunity to compare proteins with completely distinct folds, and originating from various sources and amphipathic environments (Fig. 4., STable 1., SFig. 6–11.). Within each protein, the hydrophobic belt distances cluster rather well, showing a narrow distribution of distances, sampling apparently randomly across the distribution of all proteins shown in Fig. 2. Comparison of all these proteins reveals that their means group in similar ranges, with overlapping confidence interval of the mean, invoking a comparable hydrophobic belt around all these proteins.

**Fig. 4.**
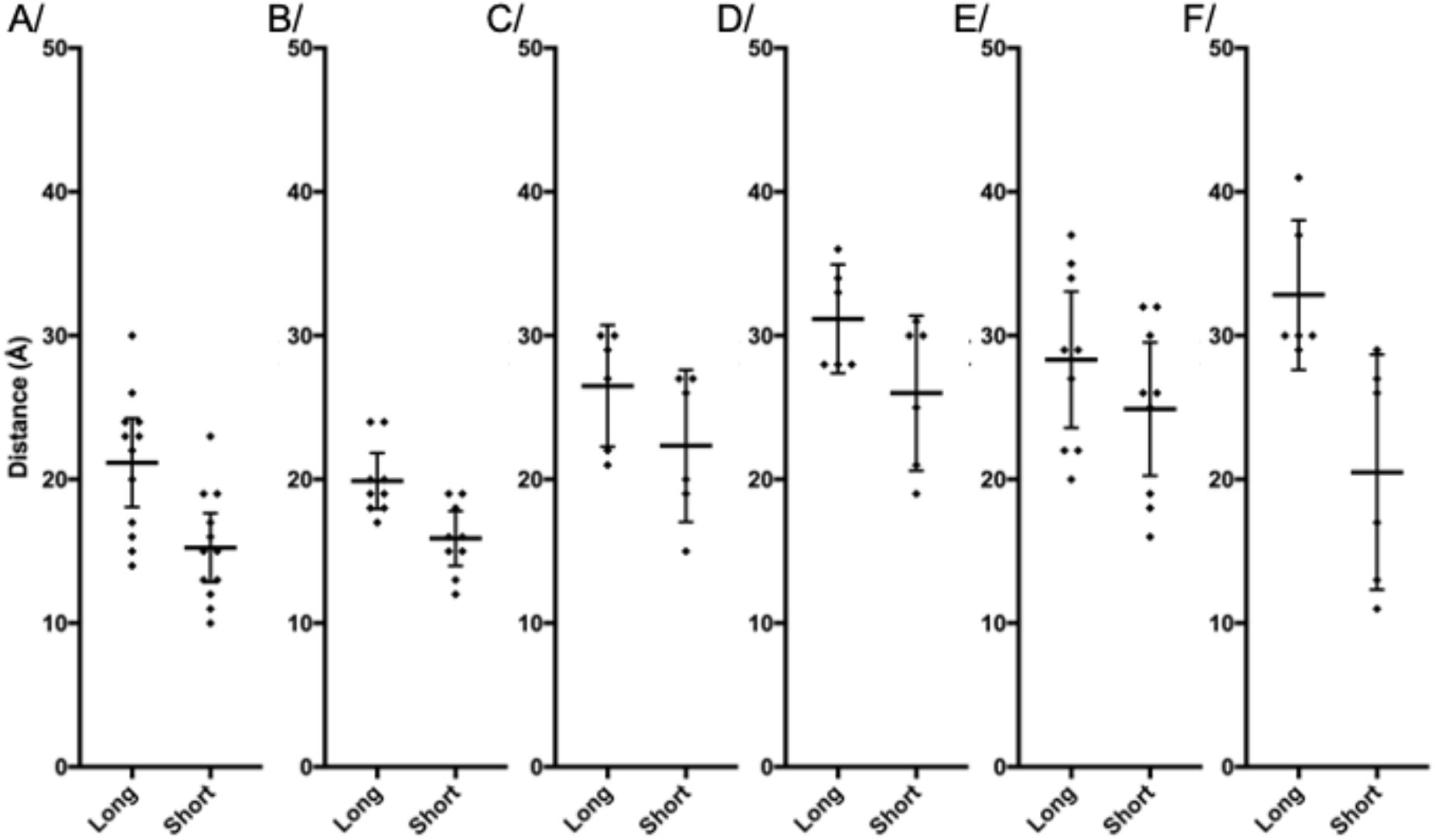
Distance distributions between the protein edge and the belt boundary for several types of proteins. Given the limited amount of structures in each family, nanodiscs, amphipols and detergents have been reported together. Each dot corresponds to a distance, the horizontal bar is the mean with the error bar displaying the 95% confidence interval of the mean (also in brackets below). Long and short distances are separated for clarity. **A/**PDK-TRP family, long 21 Å [18–24] and short 15 Å [13–18]. **B/**V-ATPase, long 20 Å [18–22] and short 16 Å [14–18]. **C/**OTP3, long 27 Å [22–31] and short 22 Å [17–28]. **D/**OSCA, long 31 Å [27–35] and short 26 Å [21–31]. **E/**LRRC8A, long 28 Å [24–33] and short 25 Å [20–30]. **F/**TMEM16, long 33 Å [28–38] and short 21 Å [12–29].

### 2.5. The superfamily of ABC transporters

ATP-binding Cassette (ABC) transporters are a large superfamily of transporters, harnessing the energy of ATP-binding and hydrolysis to translocate a wide range of substrate across many biological membranes. They are ubiquitous, and involved in many important cell-homeostasis functions[10]. While no single ABC transporter has been solved by cryo-EM in different hydrophobic environment, these proteins display a common fold and all together have been solved in nanodiscs, amphipols and detergents. They also offer the advantage that there is a large amount of structures, solved by several groups around the world using their own methodologies, and that their structures have been solved in multiple conformations offering a unique view of the solvent distribution around proteins in motion. Type I (or Type-V exporter, ABCB1-like) and type II (or Type-VI exporter, ABCG2-like) have been separated for clearer analysis (Fig. 5AB., SFig. 12–14.). Within the type I, no difference is detected between the distance distributions. Between type I and II, the distances are also inseparable, claiming that the hydrophobic belt around ABC transporters is always of similar size, regardless of the conformation or the arrangement of trans-membrane helices.

**Fig. 5.**
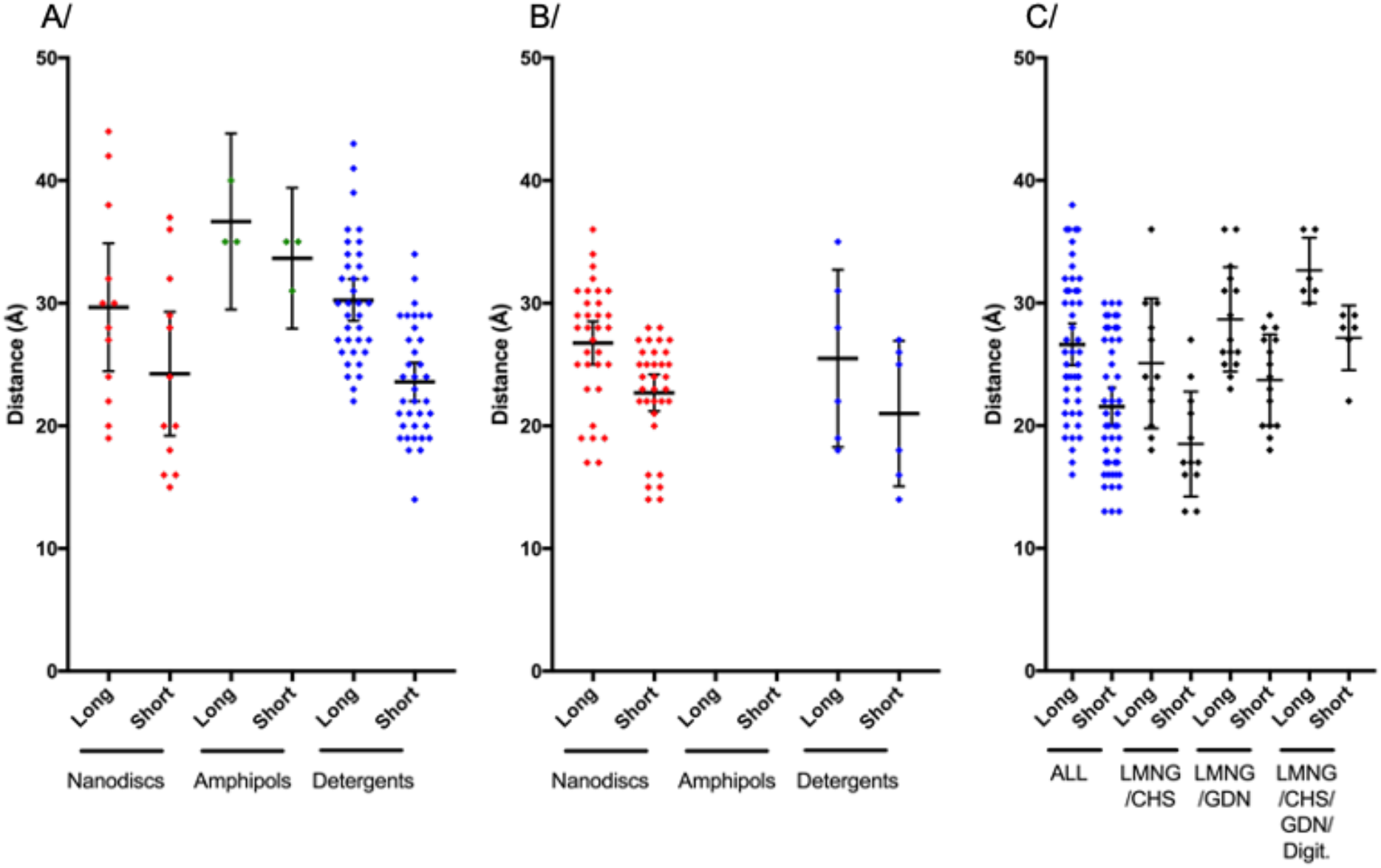
Distance distributions around ABC transporters and GPCRs. **A**/ Type V ABC transporter (ABCB1-like). Nanodiscs, long 30 Å [24–35] and short 25 Å [19–29]. Amphipols, long 37 Å [29–44] and short 34 Å [28–39]. Detergents, long 30 Å [29–32] and short 24 Å [22–25]. **B**/ Type VI ABC transporter (ABCG2-like). Nanodiscs, long 27 Å [25–28] and short 23 Å [21–24]. Detergents, long 26 Å [18–33] and short 21 Å [15–27]. **C**/ GPCR, all structures, in detergents: long 27 Å [25–28] and short 22 Å [20–23]. For structures solved, in the mixture LMNG/CHS: long 25 Å [22–29] and short 19 Å [16–21]; LMNG/GDN: long 29 Å [26–31] and short 24 Å [22–26], and for the complex mixture of LMNG/CHS/GDN with or without digitonin: long 33 Å [30–36] and short 27 Å [24–30].

### 2.6. Similar detergent belt reconstructions around GPCRs

Twenty one unique structures of GPCR were found in the present database, belonging to the A, B, C or F classes (or G, R, F or S, respectively, according to the GRAFS nomenclature[11]), all solved in detergents. The vast majority used LMNG as a base, alone or in combination with other cholesterol-like detergents such as CHS, GDN or digitonin. All structures have been solved in complex with their cognate G proteins, and/or β-arrestin, in various flavors. Like for ABC transporters, all these GPCR structures share an overall fold that grants the direct comparison of their associated belts, with local differences between structures making it more worthwhile to analyze differences in the detergent belt measurements. The detergent belt distance distribution (Fig. 5., SFig. 15–16.) is inseparable from the ABC transporter ones, or from the global distance distribution of all membrane proteins solved by cryo-EM (Fig. 2.). Following this trend, the popular detergent mixes, for these GPCR structures, between LMNG and CHS, GDN or digitonin yield similar detergent belt reconstructions, on average.

### 2.7. Different types of nanodiscs yield similar reconstructions; detergent belts are all of equivalent sizes

We checked whether a difference in distance distribution can be observed among the type of hydrophobic solvent. For instance, different flavors of Membrane Scaffold Proteins (MSP) are available to form nanodiscs, varying the length of a helical fragment within the MSP to make it longer or shorter[5]. In the current dataset, proteins have been solved with 3 types of MSP, the short MSP1D1 and its longest version MSP1E3D1 comprising 3 helical insertion. MSP2N2 is formed by the fusion of two MSP1D1. Figure 6A shows the distance distribution of the nanodisc belts sorted by nanodisc type, revealing that they are undistinguishable after reconstruction. Long and short distances of two types of nanodiscs formed by MSP1D1 and MSP2N2 follow the same distribution, with means equivalent to the mean obtained for all measurements in Fig. 2. Following this observation, distances were also separated by type of detergent to distinguish if a detergent or a detergent mixture can give rise to distinct size of belts. Distances measured from different types of detergents are all virtually indissociable, and distribute in the same range as distances observed for nanodiscs and all other measurements together (Fig. 2.).

**Fig. 6.**
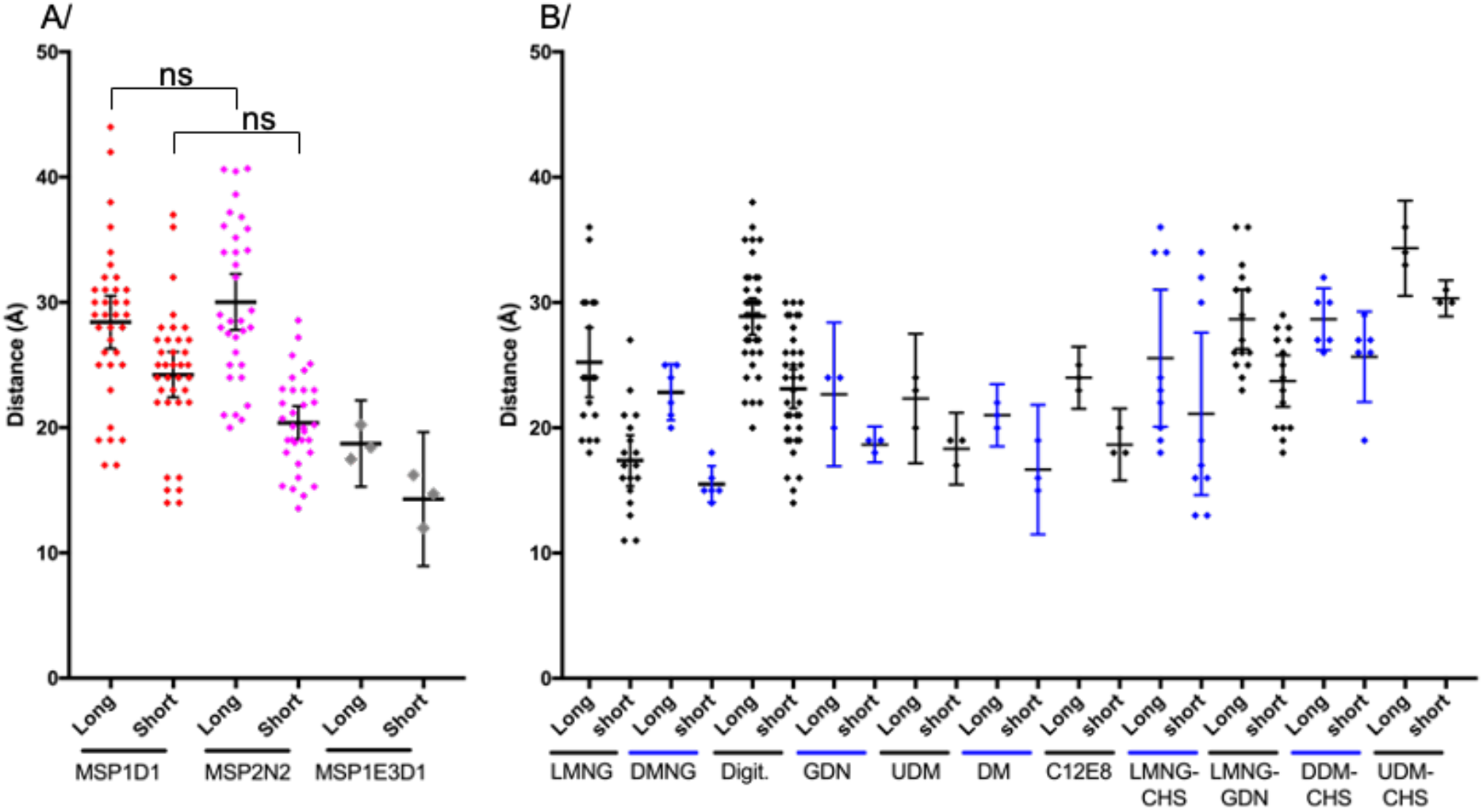
Distance distributions of the different nanodiscs or detergents belts. **A/** Distances measured for nanodiscs belts. MSP1D1, long 28 Å [26–31] and short 24 Å [22–26]. MSP2N2, long 32 Å [30–35] and short 20 Å [18–22]. MSP1E3D1, long 19 Å [15–22] and short 14 Å [9–20]. **B/** Distances measured for the most represented detergents in this dataset. LMNG (Lauryl Maltose Neopentyl Glycol), long 25 Å [22–28] and short 17 Å [15–19]. DMNG (Decyl Maltose Neopentyl Glycol), long 23 Å [20–25] and short 16 Å [14–18]. Digit. (Digitonin), long 29 Å [27–30] and short 23 Å [22–25]. GDN (Glyco-diosgenin), long 23 Å [17–28] and short 19 Å [17–20]. UDM (Undecyl-β-D-galactopyranoside), long 22 Å [17–28] and short 18 Å [15–21]. DM (Decyl-β-D-galactopyranoside), long 21 Å [19–24] and short 17 Å [12–22]. C12E8 (Octaethylene Glycol Monododecyl Ether), long 24 Å [22–27] and short 19 Å [16–22]. LMNG-CHS (CHS: Cholesteryl-hemisuccinate), long 26 Å [20–31] and short 21 Å [15–28]. LMNG-GDN, long 29 Å [26–31] and short 24 Å [22–26]. DDM-CHS (DDM: Dodecyl-β-D-galactopyranoside), long 29 Å [26–31] and short 26 Å [22–29]. UDM-CHS, long 34 Å [31–38] and short 30 Å [29–32]. Numbers are the mean followed by the 95% confidence interval of the mean in brackets.

## 3. Discussion

In order to discriminate if the compounds used to shield the membrane region of a membrane proteins have an influence on the observation of the corresponding belt by cryo-EM, we performed a statistical analysis of a curated database of selected membrane proteins solved in several hydrophobic environments. By visualizing every structure, we were able to identify in map-density distributions a signature of hydrophobic belt appearance (levels 1 & 2 in Fig. 1B.). We further identified its boundaries for every protein and measured its size for statistical analysis. 95% of all measured lengths distribute between 14 and 36 Å around the surface of trans-membrane segments, and half of the belts are comprised between 19 and 29 Å. The hydrophobic belts were further separated by type of solvent to probe whether nanodiscs, amphipols or detergents can yield tighter or larger belts. The results presented in Figure 2 show that these three types of solvent were following the same distributions, and were therefore statistically indistinguishable on average.

This result correlates well with other types of measurements of the same solvents by other methods. Molecular dynamics simulations of membrane proteins embedded in amphipols or detergents show a belt around the transmembrane regions, with some degree of flexibility[2, 12–14]. Indeed, the belt formed by these amphipathic compounds is very fluid, revealing local clusters of individual molecules, forming and deforming with time. When measured using neutron diffraction of membrane protein crystals[15], an averaging technique like cryo-EM, the detergent belt appears as a homogeneous belt around the protein. The size of the belt observed was then highly dependent on the type of crystal as the detergent could merge between belts of symmetric molecules[16]. All these techniques have been limited to the size of the system for molecular dynamic simulations, or “neutron-diffraction quality” crystals combined with deuterated detergents; here, cryo-EM allows for the visualization of any amphipathic compound, with belt measurements matching other measuring methods.

Since the hydrophobic belt has intrinsic properties to diffract electrons, it has a strong influence on the reconstructions. For example, for a 130 kDa ABC transporter, the DDM belt (400 monomers) accounts for an additional 200 kDa[2]. It is thus understandable that even if this detergent belt is not ordered, it still influences electron diffraction around the membrane protein, nicely exemplified with 3D variability of the detergent belt has been visualized in cryo-EM[17].

Nanodiscs formation with a membrane protein embedded is in itself a stunning process, where the three ingredients (Membrane Scaffold Protein, lipids and membrane protein) are mixed together, and detergents removed using biobeads. The membrane protein embedded into nanodiscs are then separated from empty nanodiscs using affinity chromatography and/or size exclusion chromatography. The object comprising the membrane protein of interest is in reality quite heterogeneous, containing a mixture of large and small nanodiscs, with more or less lipids embarked. Also, within the nanodisc, the membrane protein can move from side to side and does not always stay in the middle. This explains why the membrane scaffold protein is hardly observed in 3D reconstructions of membrane proteins in nanodiscs.

Following this idea, we further explored if we could identify within a set of protein, or type of amphipathic compound, a combination that could influence the size of the solvent belt seen around membrane proteins. We could not establish any significant difference in the measurement distributions, all falling within the overall distribution described in Figure 2. Hereabouts, the incorporation of ABC transporters and GPCRs in, this dataset yields an important viewpoint. From detergent quantification we know that the amount of detergent present around membrane proteins is directly proportional to the accessible hydrophobic area[2]. The amount of detergent around ABC transporters (12 trans-membrane helices) is thus inherently larger than the one around GPCRs (7 trans-membrane helices). One would thus expect to visualize a larger belt around ABC transporters by cryo-EM, but the size of the belt is on the contrary following the same distribution (Fig. 5.).

Finally, there is the observation that the solvent belt observed around membrane proteins by cryo-EM is circular, somewhat reminiscent of the ones observed by neutron diffraction of crystals. This is partly due to symmetries enforced during reconstructions, but at the heart, mostly due to particle averaging. Particle alignments are anchored on secondary structures, among which trans-membrane helices are a lighthouse in a fog of amphipathic solvent. The solvent observed during reconstruction is thus made out of several layers distributing radially away from the protein boundary (Fig. 7.). Level 1 corresponds to the highest density, and represents the common minimum ordered layer, where the amphipathic compound is always present around the membrane protein. This layer concomitantly increases in size and decreases in density as it radiates away from the protein boundary, representing areas of space less and less populated by the solvent. This is influenced by the fluid properties of the solvent, as the sample is vitrified in liquid ethane. Each individual particle of a dataset represents a snapshot carrying its own belt-distribution. These observations reinforce the idea that the belt visualized by cryo-EM is a fraction of the volume occupied by the hydrophobic belt and represents the common minimum ordered solvent surrounding the protein.

**Fig. 7.**
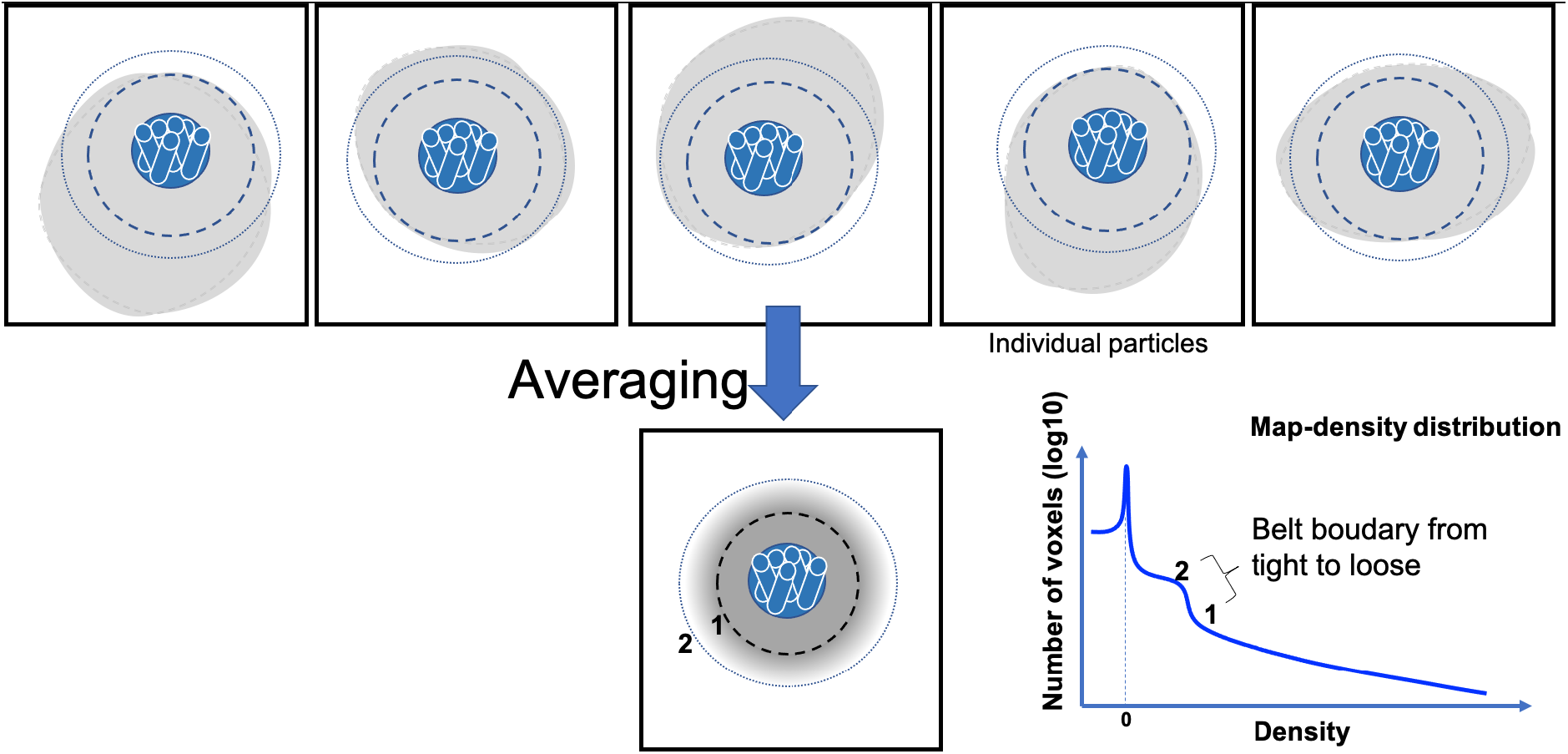
Influence of the averaging on hydrophobic solvent visualization. Top: set of particles all centered on the trans-membrane helices, with the same orientation. The inner dash circle represents the volume around the membrane protein where the hydrophobic solvent is always present. The outer dotted circle represents the spread upto where the solvent belt can be visualized. The solvent is shown in gray, with various shapes to highlight its variability around the trans-membrane domain. Bottom: The result of the averaging is a clear definition of transmembrane helices, and a gradient of presence for the solvent radiating away from the protein boundary. The level 1 and 2 correspond to the levels presented on the map-density distribution.

## 4. Material and methods

### 4.1. Membrane proteins structure database extraction

Based on the mpstruc database (https://blanco.biomol.uci.edu/mpstruc/) that lists all the membrane proteins of known 3D structure, we created a dataset containing only entries solved by Cryo-EM, as of January 17^th^, 2020. We wrote a Bash shell script in order to automatically extract information from these entries. This allowed us to determine those which have been solved in multiple hydrophobic environments (nanodiscs, amphipols or detergents) and to sort them in distinct subsets. Then, for each entry, we extracted from the Electron Microscopy Data Bank (EMDB) the map-density distribution data in order to render graphs plotting the density distribution (*i.e*. the number of voxels as a function of the density).

### 4.2. Map comparison

Maps were retrieved from EMDB and opened in ChimeraX[18]. Maps were first manually aligned, then aligned using the volume tool within ChimeraX. Threshold levels to compare the maps were adjusted to include the highest level of low contour information (level 2 in Fig 1B), without including noise voxels appearing in the box.

### 4.3. Measures of the solvent belt around the protein

The measure of solvent belt thickness was performed in ChimeraX using the tool “tape” which is included in the software. The density map histogram was used to increase or decrease the contour information. At first, the density map showed the maximum of the solvent belt information (Level 2) and vertical lines were drawn to signal the limit the solvent belt. Then the density map contour was reduced in order to see clearly the protein density (Level 0). Horizontal lines were drawn to link the vertical lines and the protein density. The tape tool measured the distance. This experiment was performed six times and in distinct positions of the solvent belt.

### 4.4. Statistical analysis

Statistical analysis (Prism.v7.2) was performed only when the amount of measures was sufficient to perform meaningful statistics. For this reason, some measures in amphipols or on individual types of proteins or hydrophobic environments were excluded. ANOVA was used to distinguish differences between means, and Kruskal-Wallis was used to distinguish differences on ranks. For all figures, means were computed as well as the 95% confidence interval of the mean.

## Author contributions

XR extracted the database. VZ measured all distances. VZ, AG and VC created the figures and analysed the data. All authors wrote the manuscript.

## Acknowledgments

This work was supported by the CNRS, Lyon University and the French National Research Agency, ANR-CLAMP2-18-CE11-0002-01 to PF and VC and ANR-19-CE11-0023-01 to VC and PF.

---

**STable 1:** List of proteins included in the study. The table relates for each protein the Protein Data Bank (PDB) entry, the Electron Microscopy Database (EMD) entry, the release year, the symmetry employed, the overall resolution, the nature of the amphipathic compound used to solve the structure (final purification step), and the reference of the structure.

| PROTEIN | PDB | EMD | YEAR | SYM. | RES.<br>(Å) | Amphipathic<br>compound | Reference |
| --- | --- | --- | --- | --- | --- | --- | --- |
| TRPV1 | 3J5P | 5778 | 2013 | C4 | 3.3 | Amphipol | [19] |
|  | 3J5Q | 5776 | 2013 | C4 | 3.8 | Amphipol | [20] |
|  | 3J5R | 5777 | 2013 | C4 | 4.2 | Amphipol | [20] |
|  | 5IRX | 8117 | 2016 | C4 | 2.95 | Nanodisc (MSP2N2) | [21] |
|  | 5IRZ | 8118 | 2016 | C4 | 3.28 | Nanodisc (MSP2N2) | [21] |
|  | 5IS0 | 8119 | 2016 | C4 | 3.4 | Nanodisc (MSP2N2) | [21] |
| TRPV2 | 6U84 | 20677 | 2019 | C4 | 3.7 | Nanodisc (MSP2N2) | [22] |
|  | 5AN8 | 6455 | 2015 | C4 | 3.8 | Amphipol | [23] |
|  | 6O03 | 20143 | 2019 | C4 | 2.9 | Nanodisc (MSP2N2) | [24] |
|  | 6B04 | 7118 | 2018 | C4 | 4.0 | LMNG | [25] |
|  | 5HI9 | 6580 | 2016 | C4 | 4.4 | DMNG | [26] |
| TRPV3 | 6DVW | 8919 | 2018 | C4 | 4.3 | Digitonin | [27] |
|  | 6DVY | 8920 | 2018 | C4 | 4.0 | Digitonin | [27] |
|  | 6DVZ | 8921 | 2018 | C4 | 4.24 | GDN | [27] |
|  | 6LGP | 882 | 2019 | C4 | 3.3 | Nanodisc (MSP2N2) | [28] |
|  | 6MHO | 9115 | 2018 | C4 | 3.4 | Poly (Maleic Anhydride-alt-1-Decene)/PMALC8 | [29] |
|  | 6UW4 | 20917 | 2020 | C4 | 3.1 | Nanodisc (MSP2N2) | [30] |
|  | 6MHS | 9117 | 2018 | C4 | 3.2 | Poly (Maleic Anhydride-alt-1-Decene)/PMALC8 | [29] |
|  | 6UW6 | 20918 | 2020 | C4 | 3.66 | Nanodisc (MSP2N2) | [30] |
|  | 6UW9 | 20920 | 2020 | C4 | 4.33 | Nanodisc (MSP2N2) | [30] |
|  | 6UW8 | 20919 | 2020 | C4 | 4.02 | Nanodisc (MSP2N2) | [30] |
|  | 6PVL | 20492 | 2019 | C4 | 4.4 | GDN | [31] |
|  | 6PVM | 20493 | 2019 | C4 | 4.5 | GDN | [31] |
|  | 6PVO | 20495 | 2019 | C4 | 5.18 | GDN | [31] |
|  | 6PVN | 20494 | 2019 | C4 | 4.07 | GDN | [31] |
|  | 6PVP | 20496 | 2019 | C4 | 4.48 | GDN | [31] |
| TRPV5 | 6B5V | 7058 | 2017 | C4 | 4.8 | DMNG | [32] |
|  | 6O1N | 593 | 2019 | C4 | 2.9 | Nanodisc (MSP2N2) | [33] |
|  | 6PBF | 20292 | 2019 | C4 | 4.2 | Nanodisc (MSP2N2) | [32] |
| TRPV6 | 6D7T | 7825 | 2018 | C4 | 4.44 | Amphipol | [34] |
|  | 6E2F | 8961 | 2018 | C1 | 3.9 | Amphipol | [34] |
|  | 6E2G | 8962 | 2018 | C1 | 3.6 | Amphipol | [34] |
| LRRC8A | 5ZSU | 6952 | 2018 | C3 | 4.25 | Digitonin | [35] |
|  | 6O00 | 564 | 2019 | C6 | 4.18 | Nanodisc (MSP1E3D1) | [36] |
|  | 6DJB | 7935 | 2018 | C3 | 4.4 | Digitonin | [37] |
| TMEM16 | 6BGI | 7095 | 2017 | C2 | 3.8 | Nanodisc (MSP2N2) | [38] |
|  | 6BGJ | 7096 | 2017 | C2 | 3.8 | LMNG | [38] |
| V-ATPase | 5TJ5 | 8409 | 2016 | C1 | 3.9 | Amphipol | [39] |
|  | 6C6L | 7348 | 2018 | C1 | 3.5 | Nanodisc (MSP1E3D1) | [40] |
|  | 6O7T | 644 | 2019 | / | 3.2 | GDN | [41] |
| OTP3 | 6NF6 | 9361 | 2019 | C2 | 3.3 | Nanodisc (MSP2N2) | [42] |
|  | 6O84 | 650 | 2019 | - | 3.92 | DDM CHS | [43] |
| OSCA | 6MGV | 9112 | 2018 | C2 | 3.1 | Nanodisc (MSP2N2) | [44] |
|  | 6OCE | 20017 | 2019 | C2 | 4.9 | UDM-CHS | [45] |
| PKD TRP | 5T4D | 8354 | 2016 | - | 3.0 | Nanodisc | [46] |
|  | 6A70 | 6991 | 2018 | - | 3.6 | Digitonin | [47] |
|  | 5K47 | 8200 | 2016 | - | 4.2 | UDM | [48] |
|  | 5MKF | 3524 | 2017 | - | 4.2 | Amphipol | [49] |
|  | 5MKE | 3523 | 2017 | - | 4.3 | Amphipol | [49] |
| MsbA | 5TTP | 8467 | 2017 | C2 | 4.8 | Nanodisc (MSP1D1) | [50] |
|  | 5TV4 | 8469 | 2017 | C1 | 4.2 | Nanodisc (MSP1D1) | [50] |
| Pgp | 6QEE | 4536 | 2019 | C1 | 3.9 | Nanodisc (MSP1D1) | [51] |
|  | 6C0V | 7325 | 2018 | C1 | 3.4 | DDM-CHS | [52] |
|  | 6FN1 | 4281 | 2018 | C1 | 3.58 | Amphipol | [53] |
|  | 6FN4 | 4282 | 2018 | C1 | 4.14 | LMNG-CHS | [53] |
|  | 6QEX | 4539 | 2019 | - | 3.6 | Nanodisc (MSP1D1) | [51] |
| CFTR | 6D3R | 7793 | 2018 | C2 | 4.30 | Digitonin | [54] |
|  | 6MSM | 9230 | 2018 | - | 3.20 | Digitonin | [55] |
|  | 5UAK | 8516 | 2017 | - | 3.87 | Digitonin | [56] |
|  | 5UAR | 8461 | 2016 | - | 3.73 | - | [57] |
|  | 6O2P | 0611 | 2019 | - | 3.3 | - | [58] |
|  | 5W81 | 8782 | 2017 | - | 3.37 | LMNG | [59] |
| MRP1 | 5UJ9 | 8559 | 2017 | - | 3.49 | Digitonin | [60] |
|  | 5AUJ | 8560 | 2017 | - | 3.34 | Digitonin | [60] |
|  | 6BHU | 7099 | 2017 | - | 3.14 | Digitonin | [61] |
| TAP1/TAP2 | 5UJ9 | 8482 | 2017 | - | 3.97 | C12E8 | [62] |
| ABCG2 | 5NJG | 3654 | 2017 | - | 3.78 | Nanodisc (MSP1D1) | [63] |
|  | 6ETI | 3953 | 2018 | - | 3.1 | Nanodisc (MSP1D1) | [64] |
|  | 6FEQ | 4246 | 2018 | - | 3.6 | Nanodisc (MSP1D1) | [64] |
|  | 6FFC | 4256 | 2018 | C2 | 3.56 | Nanodisc (MSP1D1) | [64] |
|  | 6HCO | 0196 | 2018 | C2 | 3.58 | Nanodisc (MSP1D1) | [65] |
|  | 6HZM | 0190 | 2018 | C2 | 3.09 | Nanodisc (MSP1D1) | [65] |
| ABCA1 | 5XJY | 6724 | 2017 | C1 | 4.10 | Digitonin | [66] |
| LptB2FGC | 6MI7 | 9125 | 2019 | C1 | 4.2 | Nanodisc (MSP1D1) | [67] |
|  | 6S8N | 10125 | 2019 | C1 | 3.10 | LMNG | [68] |
| GPCR | 5UZ7 | 8623 | 2017 | C1 | 4.1 | MNG/CHS | [69] |
|  | 6B3J | 7039 | 2018 | C1 | 3.3 | MNG/CHS | [70] |
|  | 6CMO | 7517 | 2018 | C1 | 4.5 | Digitonin | [71] |
|  | 6D9H | 7835 | 2018 | C1 | 3.6 | LMNG/CHS | [72] |
|  | 6E3Y | 8978 | 2018 | C1 | 3.3 | LMNG/CHS | [73] |
|  | 6G79 | 4358 | 2018 | C1 | 3.78 | DM | [74, 75] |
|  | 6N4B | 0339 | 2019 | C1 | 3 | LMNG/GDN | [76] |
|  | 6NBF | 0410 | 2019 | C1 | 3 | LMNG/GDN/CHS | [77] |
|  | 6NI3 | 9376 | 2019 | C1 | 3,8 | LMNG | [78] |
|  | 6OIJ | 20078 | 2019 | C1 | 3,3 | LMNG/GDN | [79] |
|  | 6OIK | 20079 | 2019 | C1 | 3,6 | LMNG/GDN | [79] |
|  | 6OS9 | 20180 | 2019 | C1 | 3 | LMNG/GDN | [80] |
|  | 6OT0 | 20190 | 2019 | C1 | 3,84 | MNG/GDN/<br>CHS/ Digitonin | [81] |
|  | 6OY9 | 20222 | 2019 | C1 | 3,9 | LMNG | [82] |
|  | 6PWC | 20505 | 2019 | C1 | 4,9 | Digitonin | [83] |
|  | 6QNO | 4598 | 2019 | C1 | 4,38 | / | [84] |

**STable 2:** Belt measurements for all entries listed in STable 1.

**STable2:** Belt measurements for all entries listed in STable 1.
| PROTEIN | PDB | EMD | Belt measurements (Å) |  |  |  |  |  |
| --- | --- | --- | --- | --- | --- | --- | --- | --- |
| TRPV1 | 3J5P | 5778 | 22 | 20 | 16 | 15 | 14 | 14 |
|  | 3J5Q | 5776 | 25 | 24 | 23 | 22 | 21 | 20 |
|  | 3J5R | 5777 | 29 | 22 | 24 | 21 | 19 | 18 |
|  | 5IRX | 8117 | 33 | 29 | 28 | 16 | 15 | 15 |
|  | 5IRZ | 8118 | 36 | 32 | 32 | 17 | 17 | 14 |
|  | 5IS0 | 8119 | 36 | 32 | 32 | 19 | 18 | 17 |
| TRPV2 | 6U84 | 20677 | 40 | 41 | 36 | 20 | 23 | 22 |
|  | 5AN8 | 6455 | 29 | 29 | 27 | 17 | 16 | 16 |
|  | 6O03 | 20143 | 34 | 32 | 29 | 20 | 15 | 15 |
|  | 6B04 | 7118 | 28 | 24 | 21 | 21 | 15 | 11 |
|  | 5HI9 | 6580 | 24 | 25 | 22 | 15 | 15 | 14 |
| TRPV3 | 6DVW | 8919 | 28 | 23 | 22 | 21 | 19 | 17 |
|  | 6DVY | 8920 | 29 | 25 | 20 | 19 | 15 | 14 |
|  | 6DVZ | 8921 | 28 | 27 | 25 | 24 | 23 | 21 |
|  | 6LGP | 882 | 32 | 27 | 26 | 21 | 23 | 19 |
|  | 6MHO | 9115 | 35 | 32 | 25 | 23 | 21 | 17 |
|  | 6UW4 | 20917 | 25 | 21 | 20 | 19 | 18 | 16 |
|  | 6MHS | 9117 | 24 | 20 | 19 | 18 | 17 | 15 |
|  | 6UW6 | 20918 | 34 | 33 | 28 | 24 | 23 | 22 |
|  | 6UW9 | 20920 | 28 | 24 | 21 | 20 | 19 | 18 |
|  | 6UW8 | 20919 | 26 | 25 | 24 | 23 | 23 | 19 |
|  | 6PVL | 20492 | 25 | 23 | 23 | 22 | 21 | 16 |
|  | 6PVM | 20493 | 32 | 30 | 29 | 27 | 23 | 18 |
|  | 6PVO | 20495 | 29 | 23 | 22 | 21 | 20 | 19 |
|  | 6PVN | 20494 | 31 | 24 | 23 | 21 | 18 | 15 |
|  | 6PVP | 20496 | 35 | 31 | 29 | 22 | 21 | 19 |
| TRPV5 | 6B5V | 7058 | 25 | 21 | 20 | 18 | 16 | 15 |
|  | 6O1N | 593 | 35 | 34 | 27 | 17 | 15 | 14 |
|  | 6PBF | 20292 | 37 | 39 | 36 | 25 | 22 | 21 |
| TRPV6 | 6D7T | 7825 | 20 | 19 | 17 | 15 | 15 | 13 |
|  | 6E2F | 8961 | 41 | 31 | 25 | 24 | 22 | 20 |
|  | 6E2G | 8962 | 21 | 20 | 18 | 17 | 14 | 13 |
| LRRC8A | 5ZSU | 6952 | 29 | 29 | 27 | 26 | 26 | 25 |
|  | 6O00 | 564 | 37 | 34 | 35 | 32 | 32 | 30 |
|  | 6DJB | 7935 | 22 | 22 | 20 | 19 | 18 | 16 |
| TMEM16 | 6BGI | 7095 | 41 | 37 | 29 | 29 | 27 | 26 |
|  | 6BGJ | 7096 | 30 | 30 | 30 | 17 | 13 | 11 |
| V-ATPase | 5TJ5 | 8409 | 19 | 19 | 17 | 16 | 15 | 13 |
|  | 6C6L | 7348 | 20 | 18 | 18 | 16 | 15 | 12 |
|  | 6O7T | 644 | 24 | 24 | 20 | 19 | 19 | 18 |
| OTP3 | 6NF6 | 9361 | 29 | 22 | 21 | 20 | 19 | 15 |
|  | 6O84 | 650 | 30 | 30 | 27 | 27 | 27 | 26 |
| OSCA | 6MGV | 9112 | 28 | 28 | 28 | 25 | 21 | 19 |
|  | 6OCE | 20017 | 36 | 34 | 33 | 31 | 30 | 30 |
| PKD TRP | 5T4D | 8354 | 30 | 26 | 24 | 23 | 16 | 15 |
|  | 6A70 | 6991 | 24 | 23 | 20 | 19 | 19 | 17 |
|  | 5K47 | 8200 | 23 | 22 | 16 | 15 | 12 | 11 |
|  | 5MKF | 3524 | 17 | 15 | 14 | 13 | 13 | 10 |
|  | 5MKE | 3523 | 30 | 26 | 24 | 23 | 16 | 15 |
| MsbA | 5TTP | 8467 | 28 | 24 | 22 | 20 | 20 | 16 |
|  | 5TV4 | 8469 | 27 | 20 | 19 | 18 | 16 | 15 |
| Pgp | 6QEE | 4536 | 44 | 42 | 38 | 37 | 36 | 32 |
|  | 6C0V | 7325 | 32 | 27 | 26 | 29 | 19 | 26 |
|  | 6FN1 | 4281 | 40 | 35 | 35 | 35 | 35 | 31 |
|  | 6FN4 | 4282 | 36 | 34 | 34 | 34 | 32 | 30 |
|  | 6QEX | 4539 | 32 | 30 | 30 | 29 | 28 | 24 |
| BmrA | 6R81 | 4749 | 20,9 | 19,8 | 14 | 13,7 | 13,1 | 13 |
| CFTR | 6D3R | 7793 | 30 | 28 | 27 | 25 | 19 | 19 |
|  | 6MSM | 9230 | 35 | 31 | 30 | 29 | 29 | 24 |
|  | 5UAK | 8516 | 32 | 30 | 29 | 24 | 21 | 21 |
|  | 5UAR | 8461 | 43 | 41 | 33 | 29 | 28 | 25 |
|  | 6O2P | 0611 | 39 | 33 | 28 | 29 | 27 | 22 |
|  | 5W81 | 8782 | 36 | 35 | 30 | 27 | 23 | 19 |
| MRP1 | 5UJ9 | 8559 | 32 | 27 | 26 | 24 | 22 | 22 |
|  | 5AUJ | 8560 | 29 | 25 | 22 | 20 | 20 | 19 |
|  | 6BHU | 7099 | 27 | 26 | 24 | 21 | 21 | 14 |
| TAP1/TAP2 | 5UJ9 | 8482 | 25 | 24 | 23 | 20 | 18 | 18 |
| ABCG2 | 5NJG | 3654 | 20 | 19 | 17 | 16 | 15 | 15 |
|  | 6ETI | 3953 | 19 | 19 | 17 | 16 | 14 | 14 |
|  | 6FEQ | 4246 | 31 | 31 | 27 | 25 | 22 | 22 |
|  | 6FFC | 4256 | 29 | 28 | 26 | 25 | 25 | 24 |
|  | 6HCO | 196 | 33 | 32 | 31 | 28 | 28 | 26 |
|  | 6HZM | 190 | 26 | 25 | 23 | 23 | 22 | 22 |
| ABCA1 | 5XJY | 6724 | 35 | 31 | 28 | 27 | 26 | 25 |
| LptB2FGC | 6MI7 | 9125 | 28 | 25 | 23 | 22 | 21 | 20 |
|  | 6S8N | 10125 | 22 | 19 | 18 | 18 | 16 | 14 |
| GPCR | 5UZ7 | 8623 | 36 | 30 | 28 | 27 | 22 | 21 |
|  | 6B3J | 7039 | 30 | 27 | 24 | 24 | 17 | 17 |
|  | 6CMO | 7517 | 35 | 32 | 32 | 30 | 30 | 28 |
|  | 6D9H | 7835 | 23 | 22 | 19 | 19 | 17 | 16 |
|  | 6E3Y | 8978 | 24 | 20 | 18 | 16 | 13 | 13 |
|  | 6G79 | 4358 | 22 | 21 | 20 | 19 | 16 | 15 |
|  | 6N4B | 0339 | 29 | 27 | 26 | 26 | 25 | 24 |
|  | 6NBF | 0410 | 36 | 36 | 32 | 29 | 28 | 27 |
|  | 6NI3 | 9376 | 21 | 19 | 19 | 18 | 17 | 16 |
|  | 6OIJ | 20078 | 26 | 26 | 23 | 20 | 20 | 19 |
|  | 6OIK | 20079 | 33 | 31 | 31 | 28 | 27 | 22 |
|  | 6OS9 | 20180 | 25 | 25 | 24 | 23 | 20 | 18 |
|  | 6OT0 | 20190 | 31 | 31 | 30 | 29 | 28 | 22 |
|  | 6OY9 | 20222 | 24 | 24 | 24 | 21 | 20 | 16 |
|  | 6PWC | 20505 | 38 | 36 | 34 | 30 | 29 | 29 |
|  | 6QNO | 4598 | 21 | 17 | 16 | 15 | 15 | 13 |

## Supplementary Figures

**SFigure 1.**
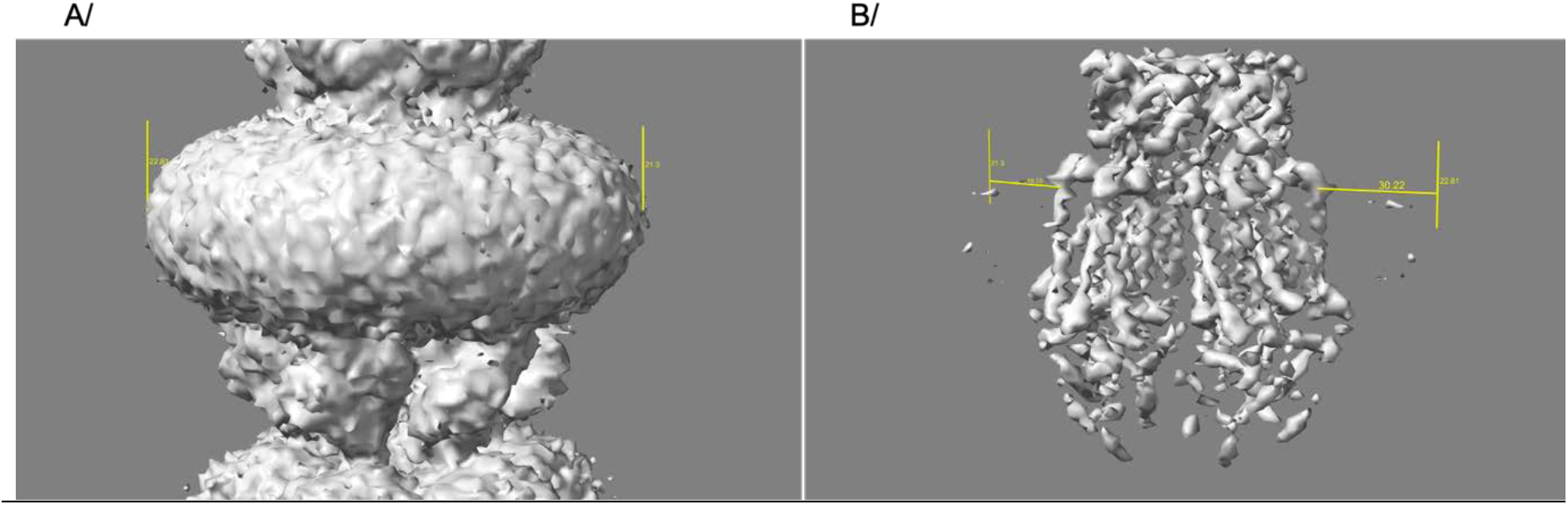
Measurement of the hydrophobic solvent size. A/ the level 2 of the hydrophobic solvent for EMD-0564 was identified (vertical yellow mark). B/ the map contour level was decreased just below level 1 to visualize the protein edge. Distances were measured using ChimeraX.

**SFigure 2.**
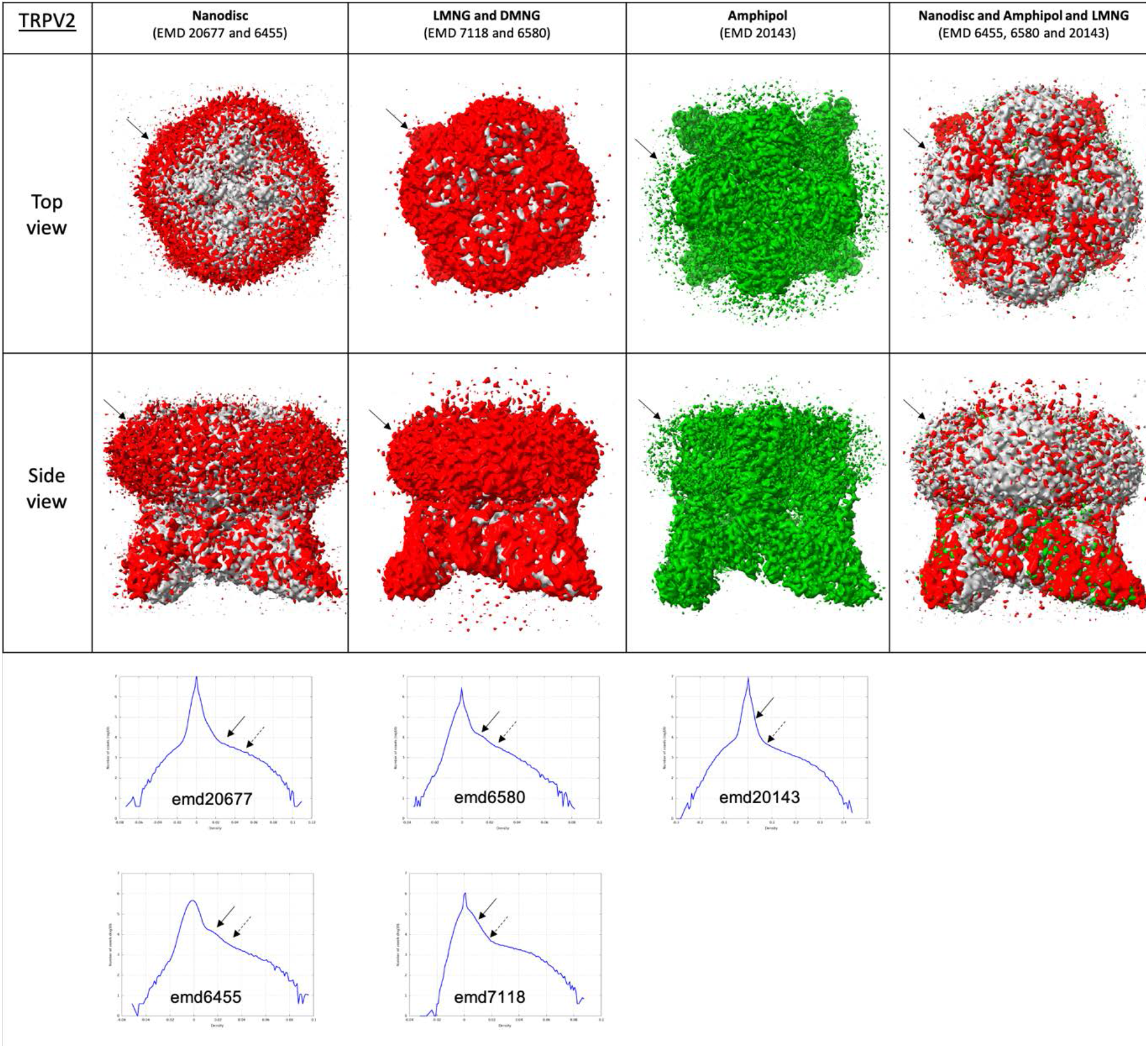
TRPV2 proteins. The top chart displays top and side views of the protein in various amphipathic environments. The last row represents a superposition of the protein solved in these environments. EMD codes are listed for reference. Map-density distributions are shown below. Solid arrows depict the amphipathic belt at level 2. Dotted arrows correspond to level 1.

**SFigure 3.**
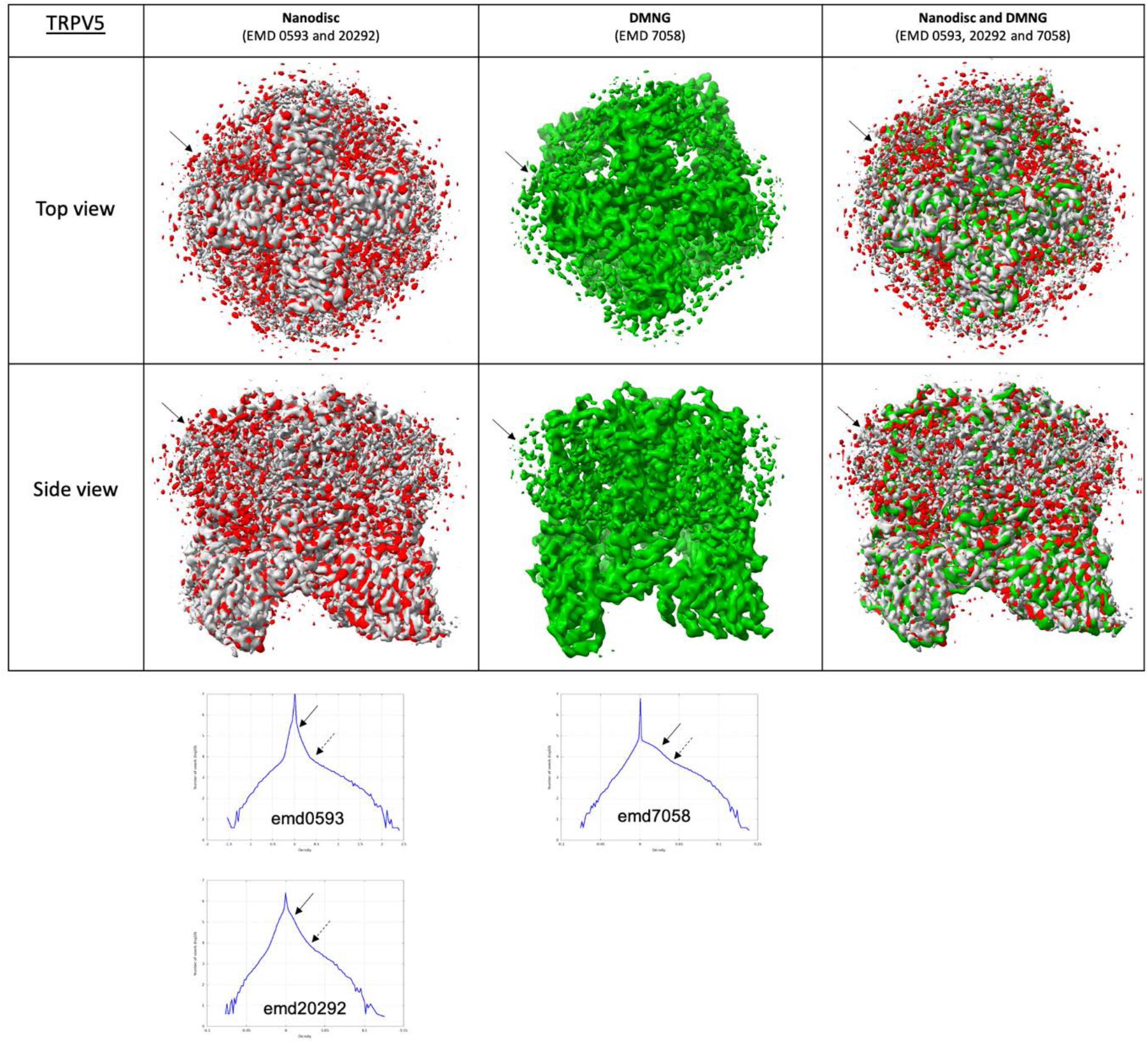
TRPV5 proteins. The top chart displays top and side views of the protein in various amphipathic environments. The last row represents a superposition of the protein solved in these environments. EMD codes are listed for reference. Map-density distributions are shown below. Solid arrows depict the amphipathic belt at level 2. Dotted arrows correspond to level 1.

**SFigure 4.**
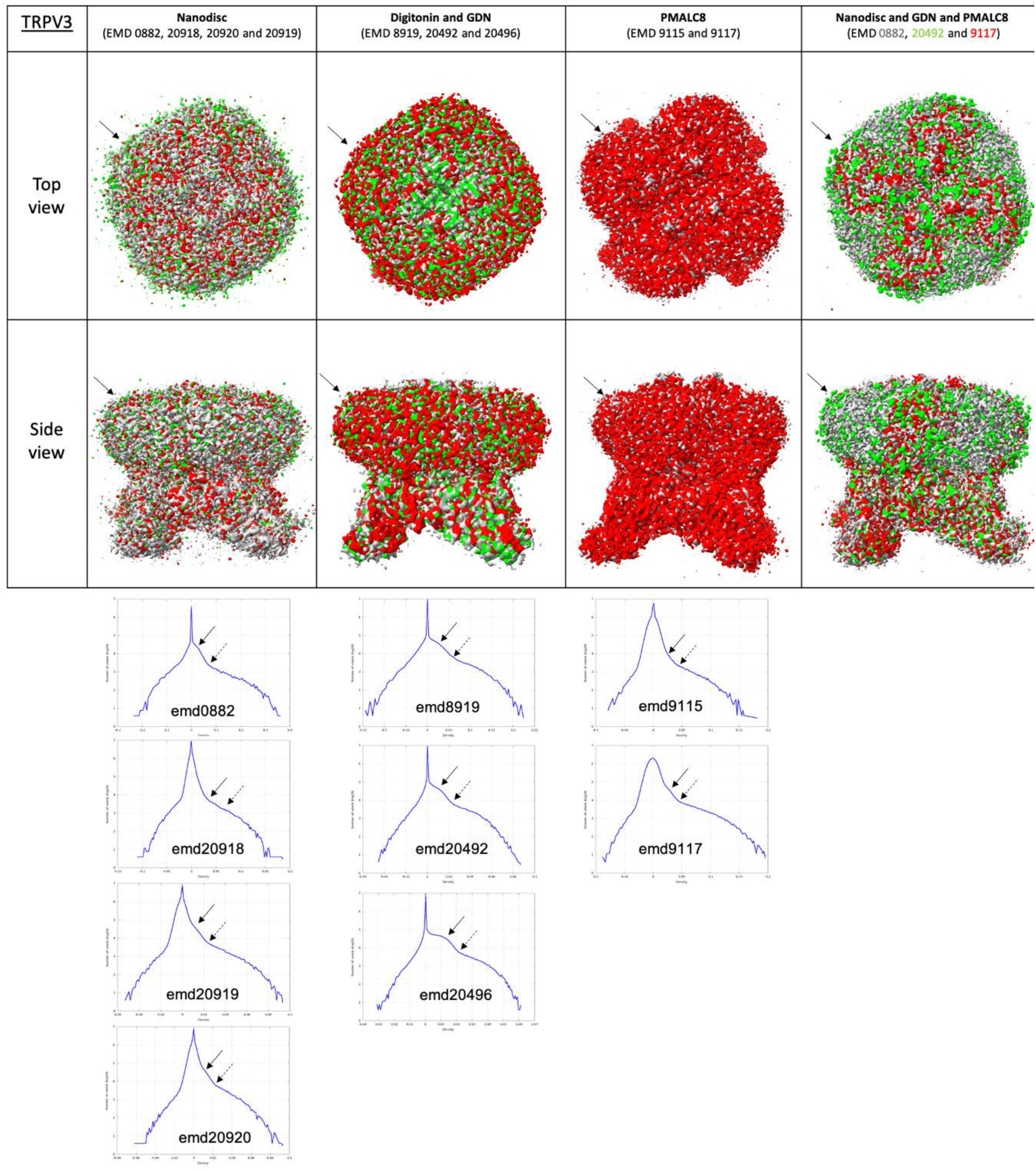
TRPV3 proteins. The top chart displays top and side views of the protein in various amphipathic environments. The last row represents a superposition of the protein solved in these environments. EMD codes are listed for reference. Map-density distributions are shown below. Solid arrows depict the amphipathic belt at level 2. Dotted arrows correspond to level 1.

**SFigure 5.**
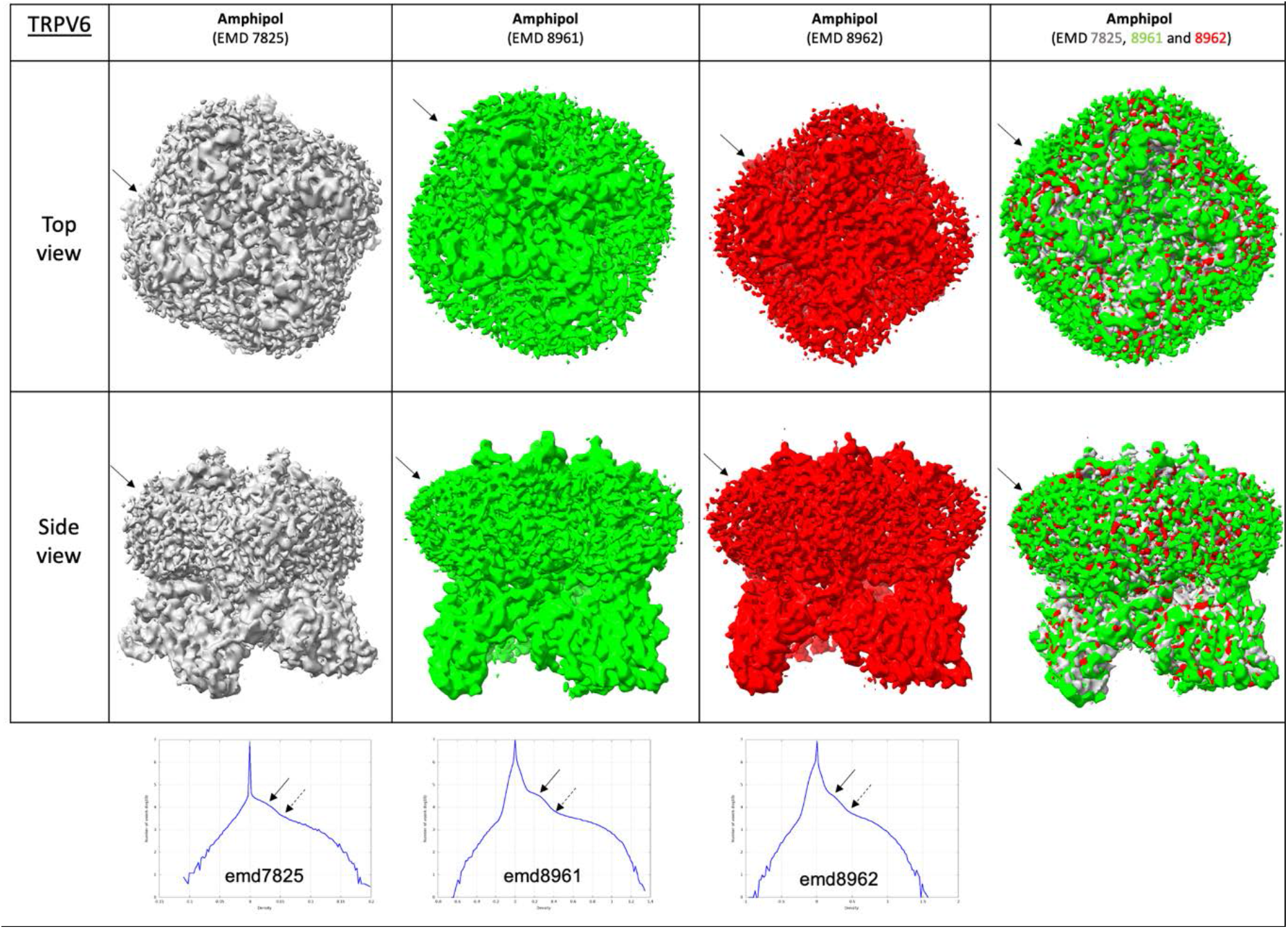
TRPV6 proteins. The top chart displays top and side views of the protein in various amphipathic environments. The last row represents a superposition of the protein solved in these environments. EMD codes are listed for reference. Map-density distributions are shown below. Solid arrows depict the amphipathic belt at level 2. Dotted arrows correspond to level 1.

**SFigure 6.**
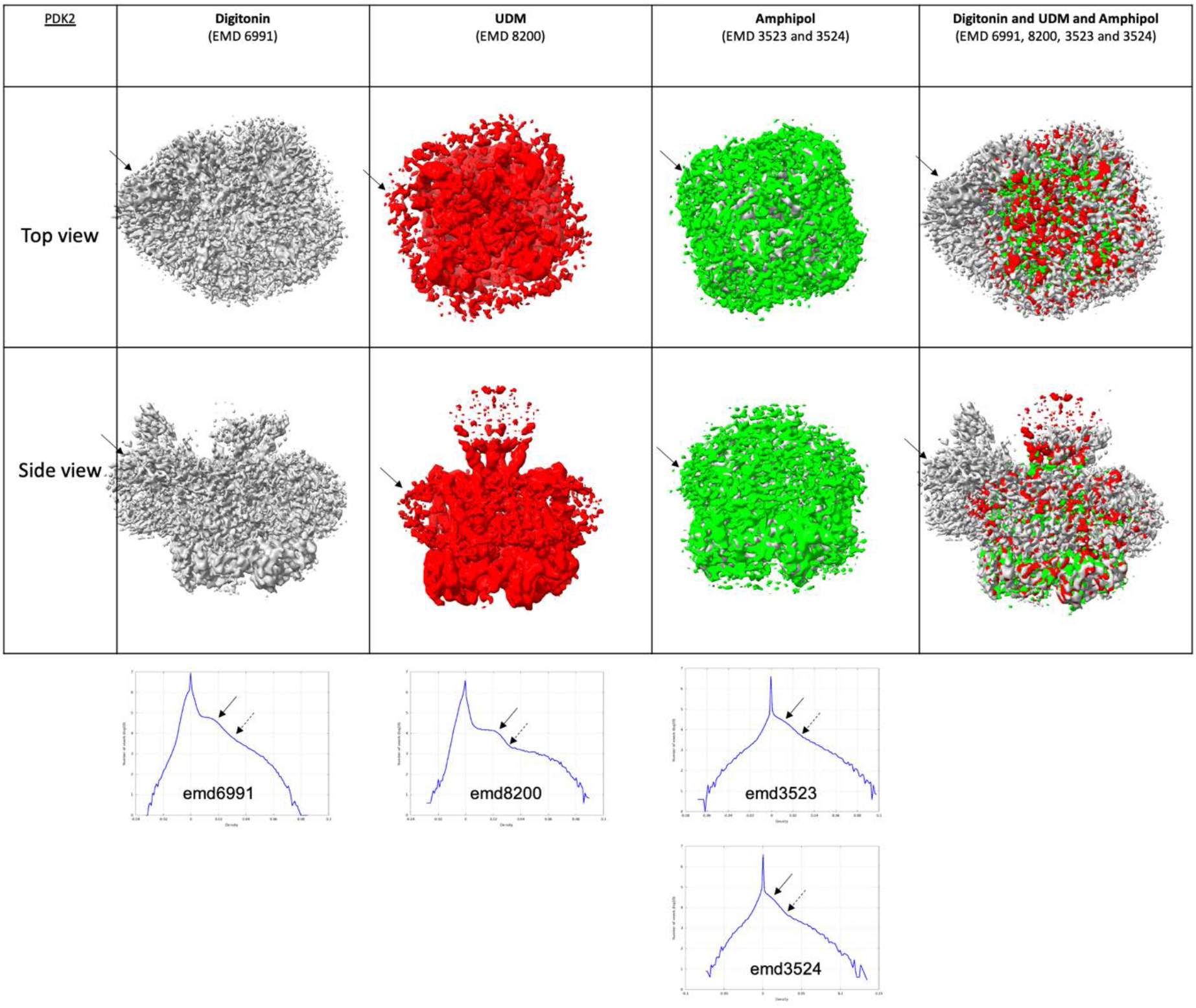
PDK2 proteins. The top chart displays top and side views of the protein in various amphipathic environments. The last row represents a superposition of the protein solved in these environments. EMD codes are listed for reference. Map-density distributions are shown below. Solid arrows depict the amphipathic belt at level 2. Dotted arrows correspond to level 1.

**SFigure 7.**
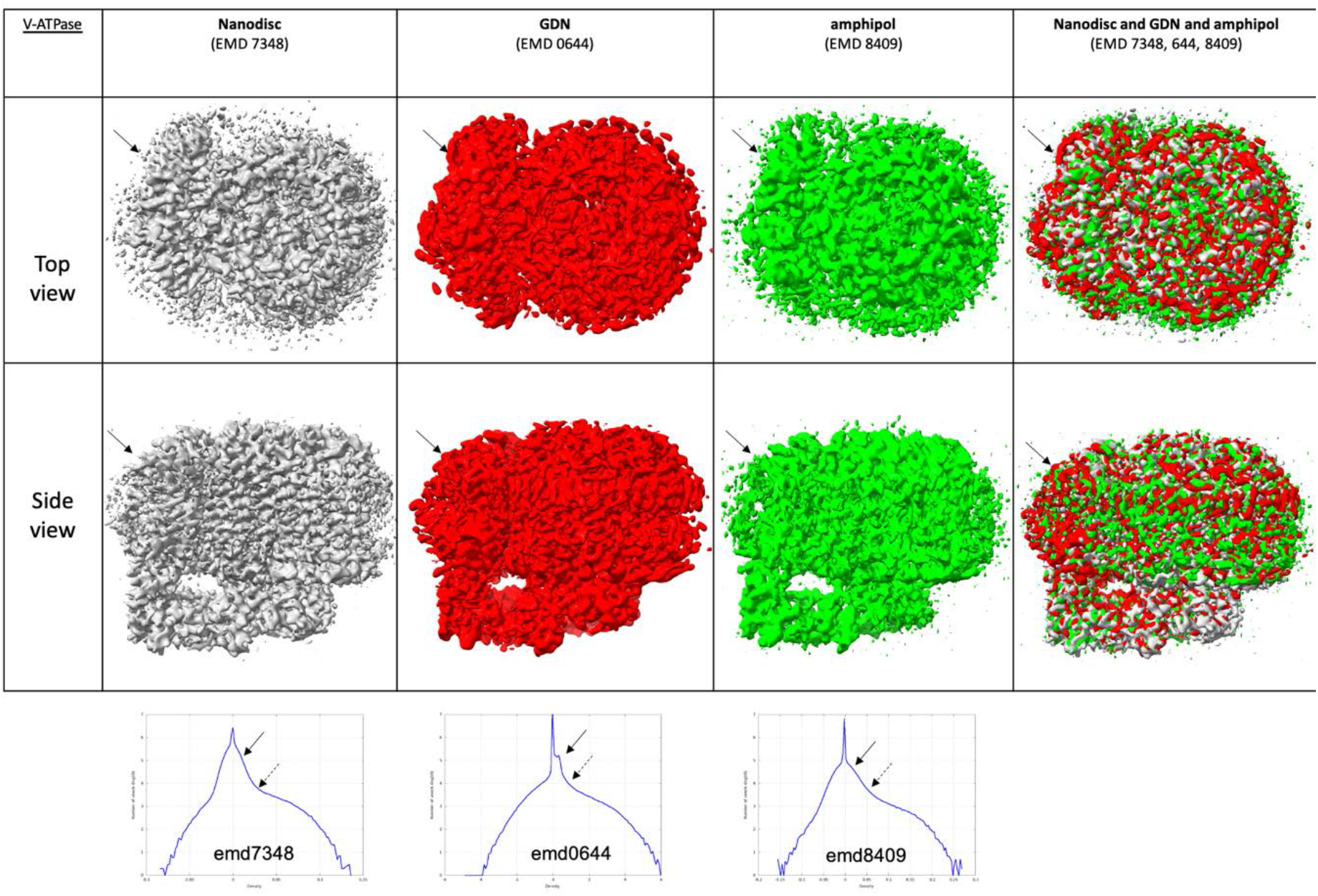
V-ATPase proteins. The top chart displays top and side views of the protein in various amphipathic environments. The last row represents a superposition of the protein solved in these environments. EMD codes are listed for reference. Map-density distributions are shown below. Solid arrows depict the amphipathic belt at level 2. Dotted arrows correspond to level 1.

**SFigure 8.**
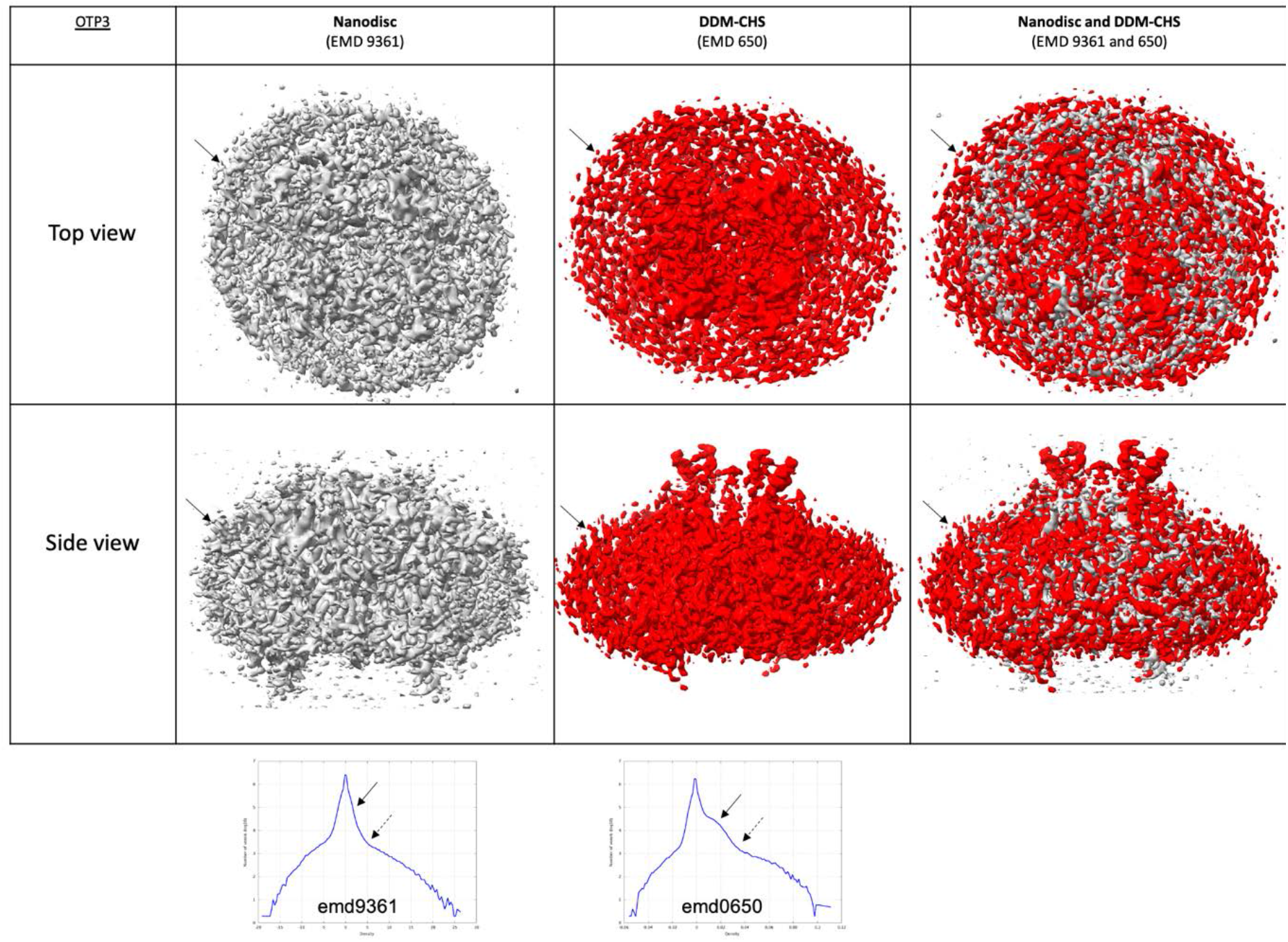
OATP3 proteins. The top chart displays top and side views of the protein in various amphipathic environments. The last row represents a superposition of the protein solved in these environments. EMD codes are listed for reference. Map-density distributions are shown below. Solid arrows depict the amphipathic belt at level 2. Dotted arrows correspond to level 1.

**SFigure 9.**
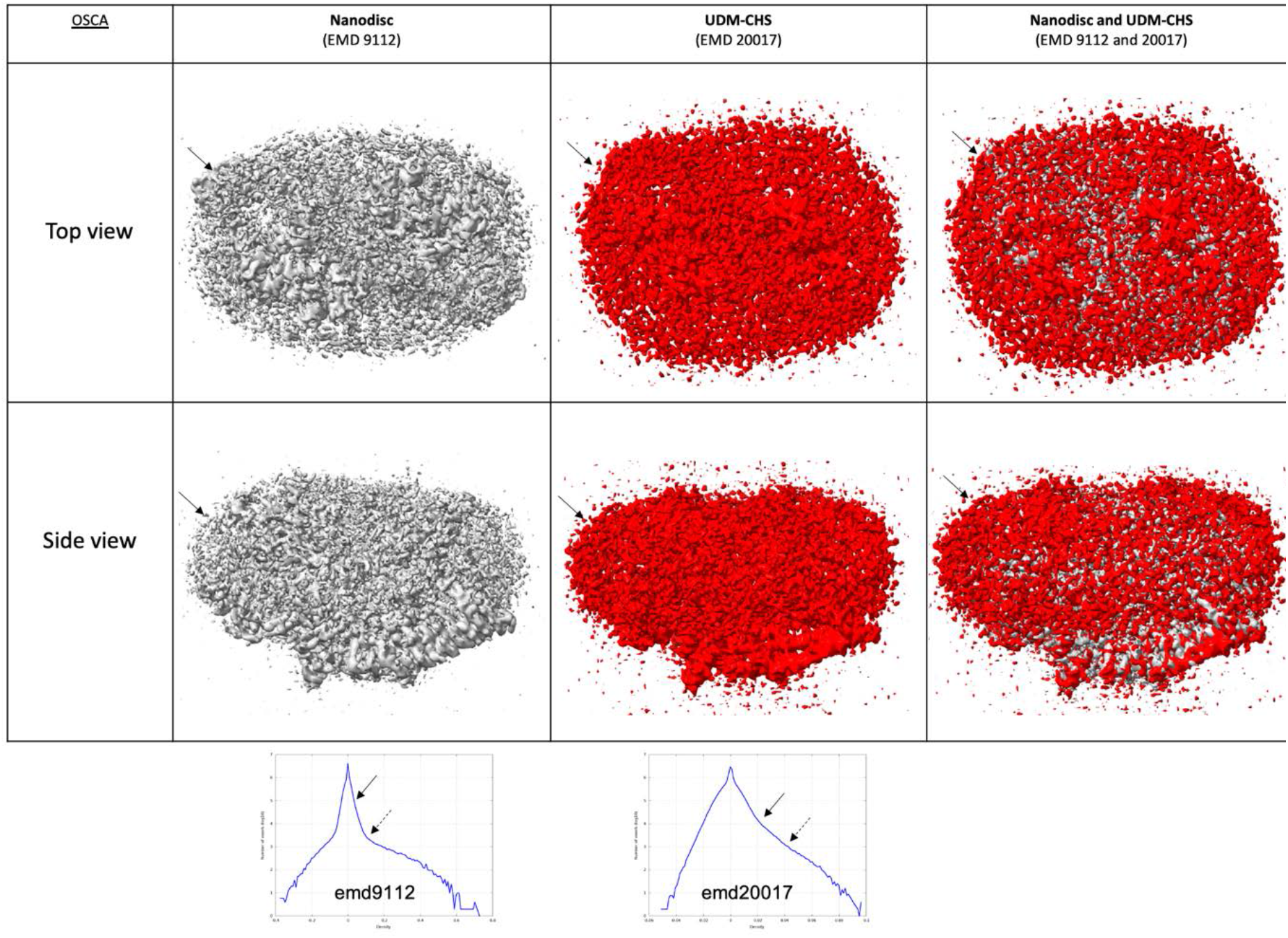
OSCA proteins. The top chart displays top and side views of the protein in various amphipathic environments. The last row represents a superposition of the protein solved in these environments. EMD codes are listed for reference. Map-density distributions are, shown below. Solid arrows depict the amphipathic belt at level 2. Dotted arrows correspond to level 1.

**SFigure 10.**
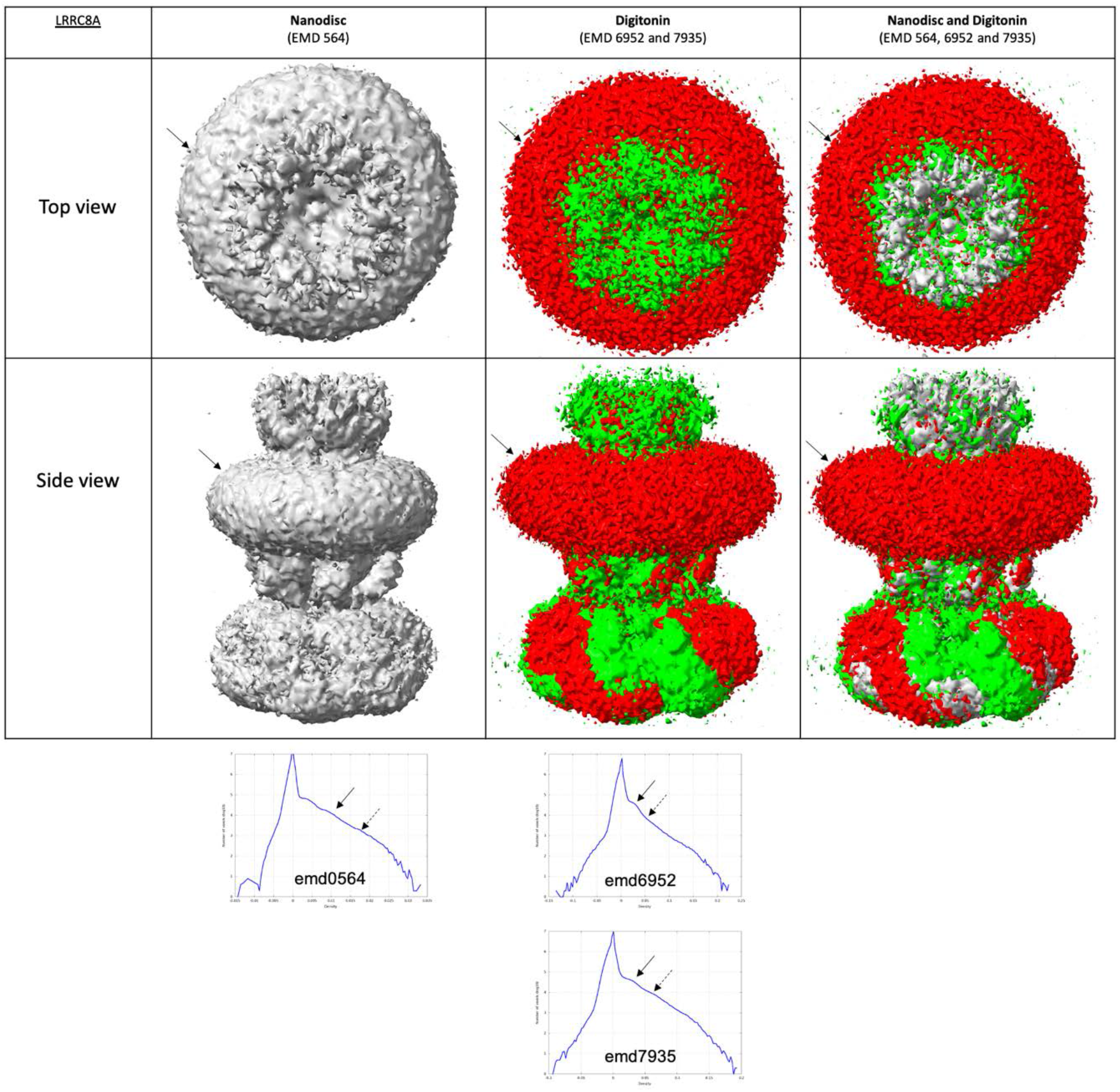
LRRC8A proteins. The top chart displays top and side views of the protein in various amphipathic environments. The last row represents a superposition of the protein solved in these environments. EMD codes are listed for reference. Map-density distributions are shown below. Solid arrows depict the amphipathic belt at level 2. Dotted arrows correspond to level 1.

**SFigure 11.**
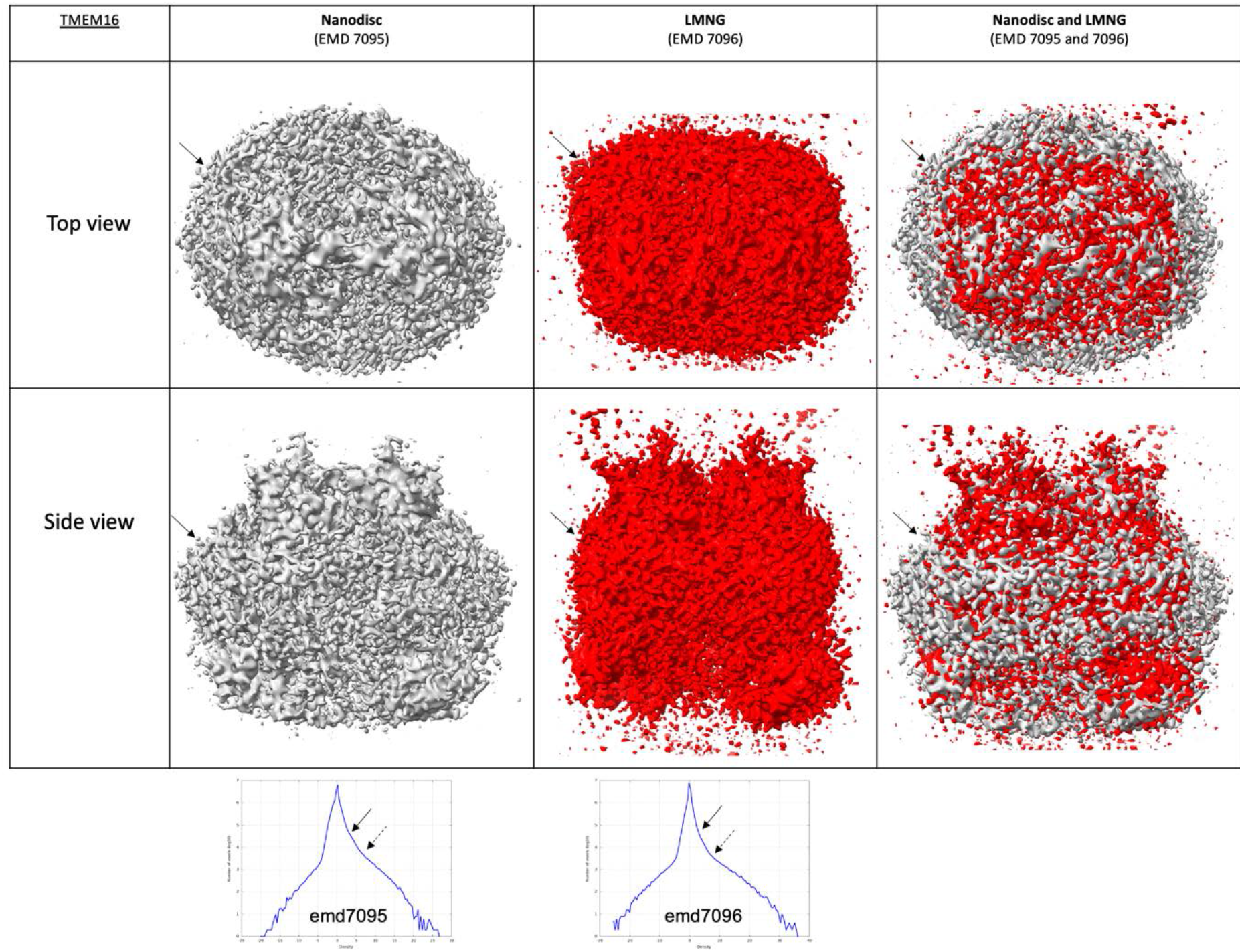
TMEM16 proteins. The top chart displays top and side views of the protein in various amphipathic environments. The last row represents a superposition of the protein solved in these environments. EMD codes are listed for reference. Map-density distributions are shown below. Solid arrows depict the amphipathic belt at level 2. Dotted, arrows correspond to level 1.

**SFigure 12.**
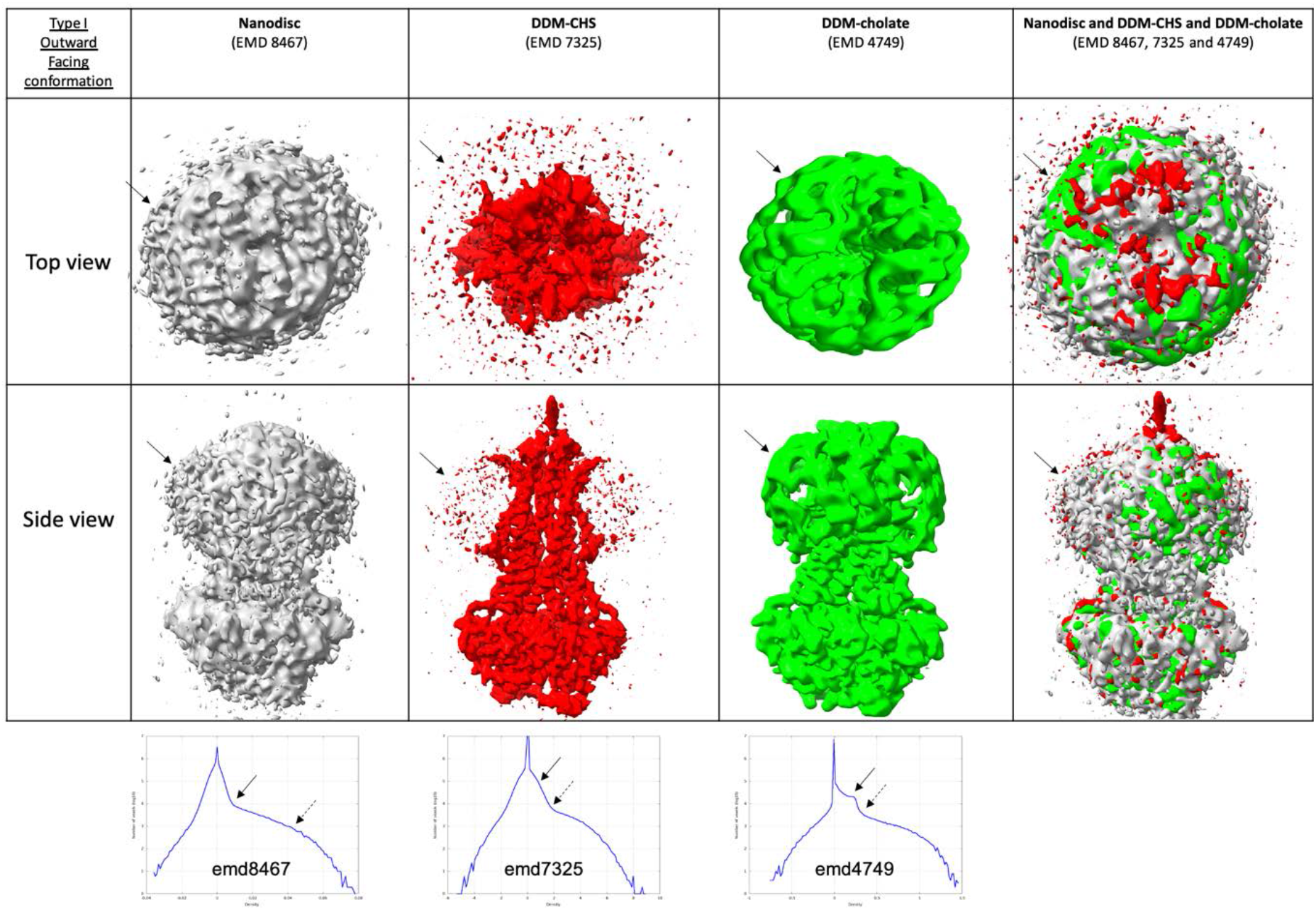
Type I ABC exporter, Outward-facing conformation. The top chart displays top, and side views of the protein in various amphipathic environments. The last row represents a superposition of the protein solved in these environments. EMD codes are listed for reference. Map-density distributions are shown below. Solid arrows depict the amphipathic belt at level 2. Dotted arrows correspond to level 1.

**SFigure 13.**
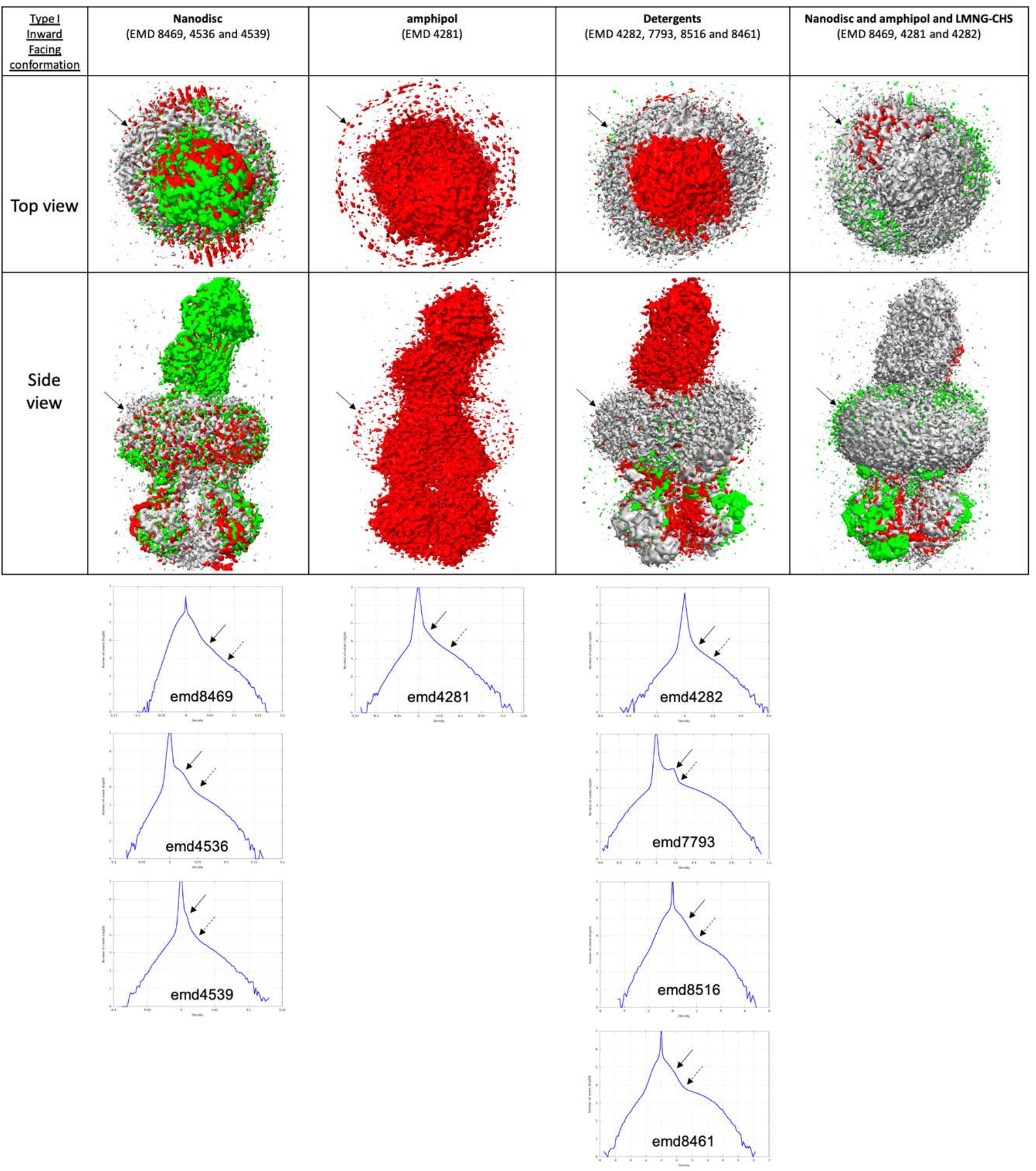
Type I ABC exporter, Inward-facing conformation. The top chart displays top and side views of the protein in various amphipathic environments. The last row represents a superposition of the protein solved in these environments. EMD codes are listed for reference. Map-density distributions are shown below. Solid arrows depict the amphipathic belt at level 2. Dotted arrows correspond to level 1. Only some proteins were displayed for clarity. All distances are available in STable 2.

**SFigure 14.**
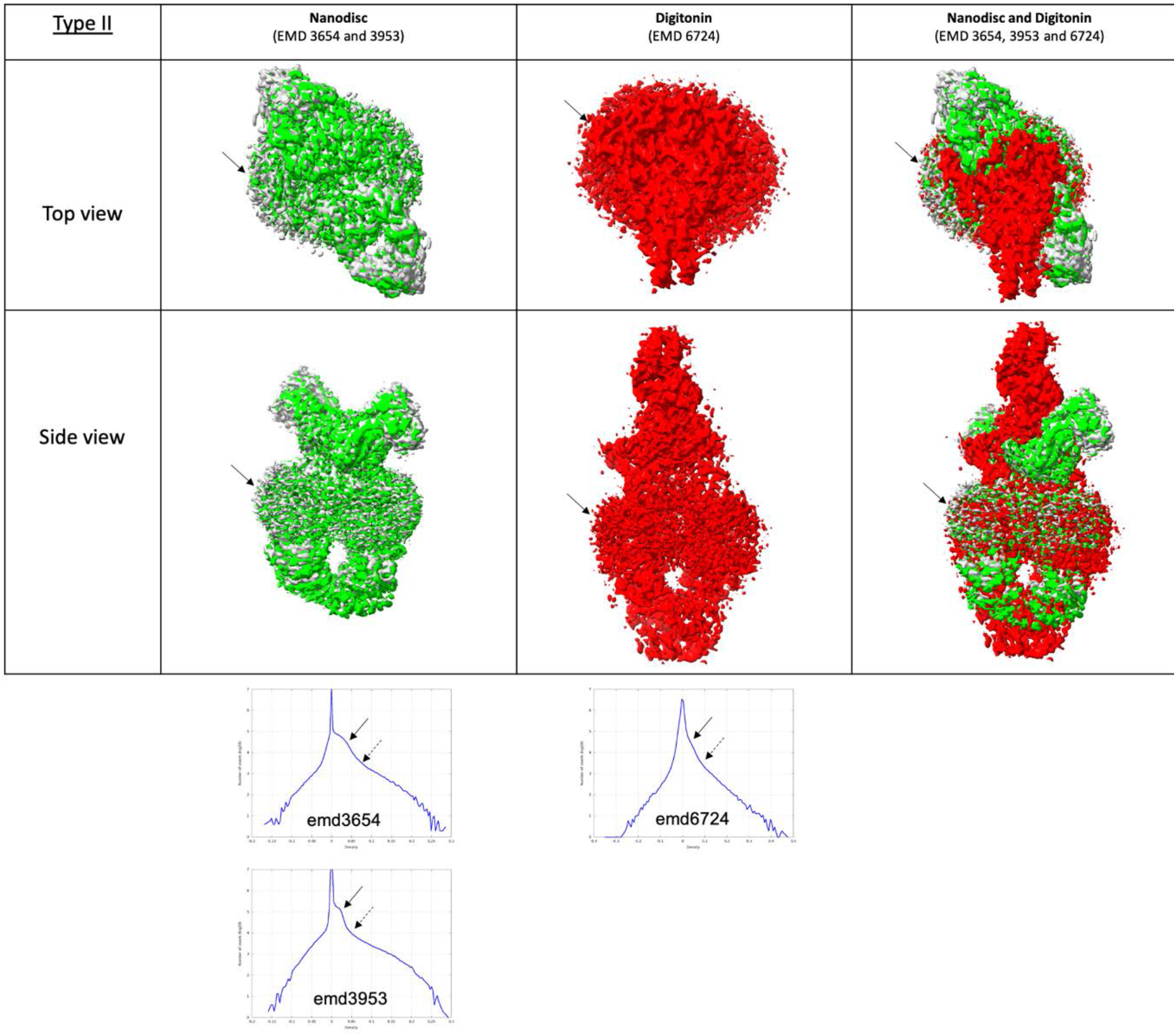
Type II ABC exporter. The top chart displays top and side views of the protein in various amphipathic environments. The last row represents a superposition of the protein solved in these environments. EMD codes are listed for reference. Map-density distributions are shown below. Solid arrows depict the amphipathic belt at level 2. Dotted arrows correspond to level 1. Only some proteins were displayed for clarity. All distances are available in STable 2.

**SFigure 15.**
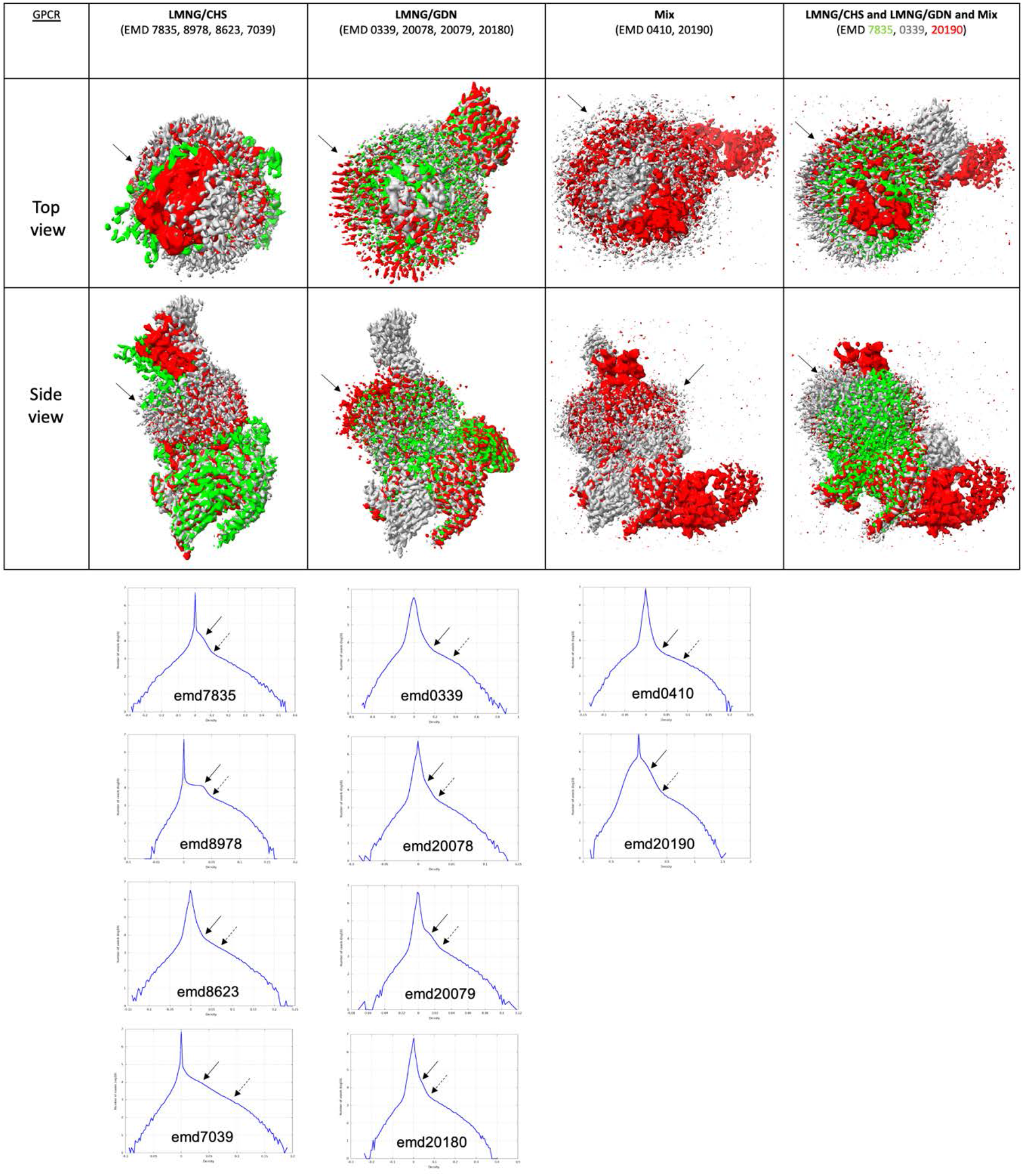
GPCR proteins (1). The top chart displays top and side views of the protein in various amphipathic environments. The last row represents a superposition of the protein solved in these environments. EMD codes are listed for reference. Map-density distributions are shown below. Solid arrows depict the amphipathic belt at level 2. Dotted arrows correspond to level 1. Only some proteins were displayed for clarity. All distances are available in STable 2.

**SFigure 16.**
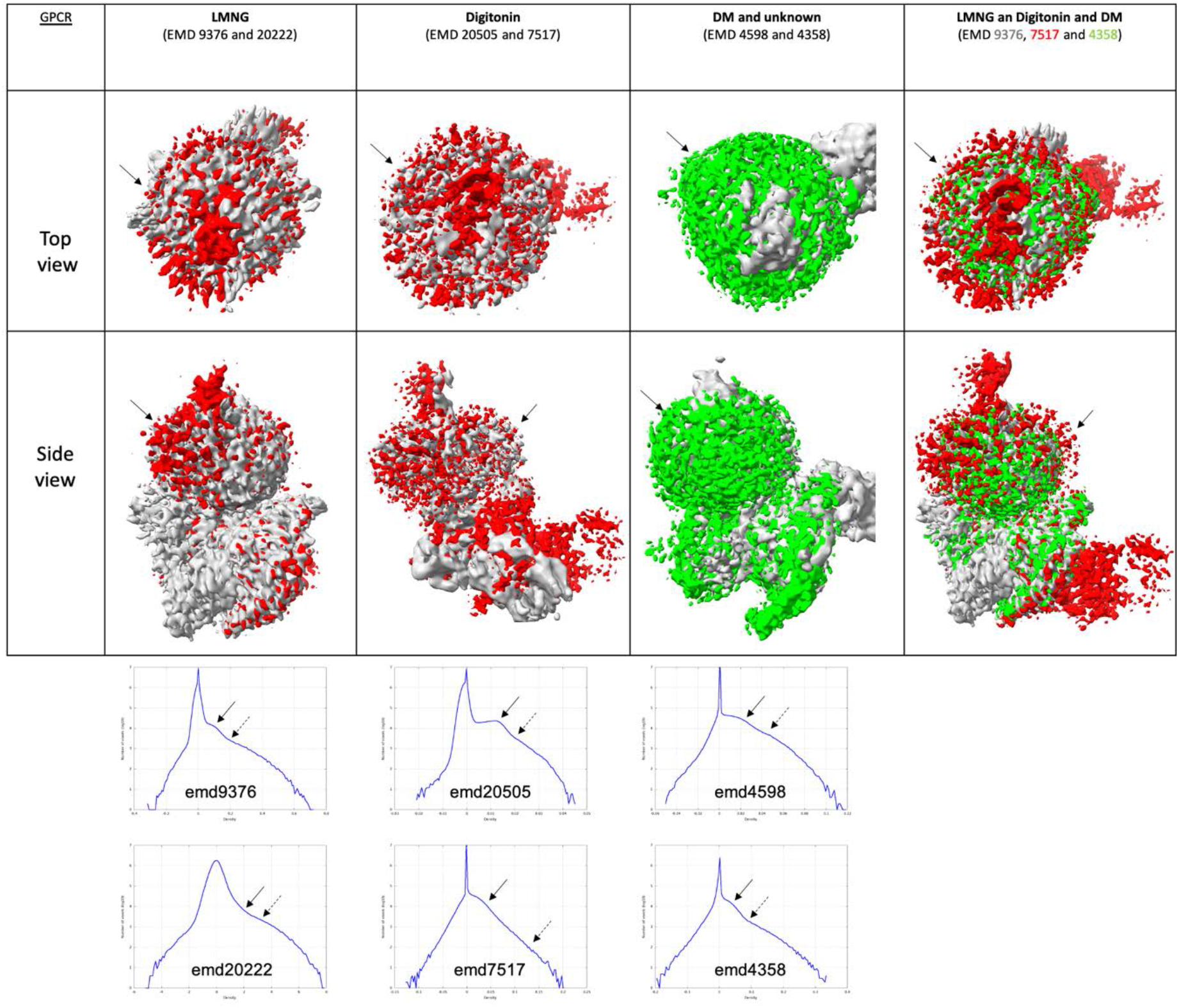
GPCR proteins (2). The top chart displays top and side views of the protein in various amphipathic environments. The last row represents a superposition of the protein solved in these environments. EMD codes are listed for reference. Map-density distributions are shown below. Solid arrows depict the amphipathic belt at level 2. Dotted arrows correspond to level 1. Only some proteins were displayed for clarity. All distances are available in STable 2.

